# Polygenic architecture of human neuroanatomical diversity

**DOI:** 10.1101/592337

**Authors:** Anne Biton, Nicolas Traut, Jean-Baptiste Poline, Benjamin S. Aribisala, Mark E. Bastin, Robin Bülow, Simon R. Cox, Ian J. Deary, Masaki Fukunaga, Hans J. Grabe, Saskia Hagenaars, Ryota Hashimoto, Masataka Kikuchi, Susana Muñoz Maniega, Matthias Nauck, Natalie A. Royle, Alexander Teumer, Maria Valdes Hernandez, Uwe Völker, Joanna M. Wardlaw, Katharina Wittfeld, Hidenaga Yamamori, Alzheimer’s Disease Neuroimaging Initiative, Thomas Bourgeron, Roberto Toro

## Abstract

We analysed the genomic architecture of neuroanatomical diversity using magnetic resonance imaging and single nucleotide polymorphism (SNP) data from >26,000 individuals from the UK Biobank project and 5 other projects that had previously participated in the ENIGMA consortium. Our results confirm the polygenic architecture of neuroanatomical diversity, with SNPs capturing from 40% to 54% of regional brain volume variance. Chromosomal length correlated with the amount of phenotypic variance captured, r∼0.64 on average, suggesting that at a global scale causal variants are homogeneously distributed across the genome. At a local scale, SNPs within genes (∼51%) captured ∼1.5 times more genetic variance than the rest; and SNPs with low minor allele frequency (MAF) captured less variance than the rest: the 40% of SNPs with MAF<5% captured <1/4th of the genetic variance. We also observed extensive pleiotropy across regions, with an average genetic correlation of r_G_∼0.45. Genetic correlations were similar to phenotypic and environmental correlations, however, genetic correlations were often larger than phenotypic correlations for the left/right volumes of the same region. The heritability of differences in left/right volumes was generally not statistically significant, suggesting an important influence of environmental causes in the variability of brain asymmetry. Our code is available at https://github.com/neuroanatomy/genomic-architecture.

## Introduction

Imaging genetics studies have traditionally emphasised the role of candidate genes and specific loci. The underlying hypothesis is that phenotypic diversity results from the action of a reduced number of genes, close to the Mendelian paradigm where a phenotype is determined by a single locus. However, the amount of neuroanatomical variance captured by candidate genes or genome-wide significant loci has remained extremely small. Genome-wide association studies (GWAS) aiming at identifying associated variants through international collaborative efforts such as ENIGMA and CHARGE have revealed only few statistically significant associated loci (Stein et al. 2012, Hibar et al. 2015, Satizabal et al. 2017), capturing <1% of phenotypic variance. This is in contrast with the large heritability of neuroanatomical diversity estimated by twin and extended pedigree studies (Blokland et al. 2012, Wen et al. 2016) which show that about 80% of neuroanatomical variability is captured by genetic factors.

The genetic architecture of neuroanatomical diversity could result, alternatively, from the aggregated effect of thousands of different loci spread across the genome, a paradigm closer to the infinitesimal model proposed by Fisher (1918). Thanks to the development of methods to estimate heritability from whole-genome genotyping data (reviewed by Yang et al. 2017), several groups have estimated that genotyped variants, taken together, were able to capture up to 55% of the phenotypic variance, retrieving about two-thirds of the heritability estimated by family studies (Toro et al. 2015, Ge et al. 2015, 2016, Zhao et al. 2018, Elliott et al. 2018). These results are compatible with the hypothesis of a highly polygenic architecture, where phenotypes are influenced by large numbers of loci of effect sizes too small to reach genome-wide significance (usually p<5×10^−8^). While information about the function of a few candidate genes can have a strong explanatory power in the case of a few causal loci, a polygenic approach calls for alternative, system-wide, sources of biological insight.

Our aim was to better understand the role of polygenic causes in the determination of neuroanatomical diversity. In a previous work, we used a sample of N = 2,011 subjects with whole-genome genotyping data from the IMAGEN consortium combined with structural magnetic resonance imaging (MRI) data analysed according to the pipelines of the ENIGMA consortium, and showed that genomic complex trait analysis (GCTA) was able to capture a substantial proportion of the variability in regional brain volume – up to 55% (Toro et al. 2015). Due to sample size limitations, however, standard errors were high (about ± 25%) and statistical power for post-hoc analyses was limited. The advent of collaborative efforts such as the ENIGMA consortium, and large scale projects such as UK Biobank, in particular, has allowed researchers to greatly increase the number of subjects used for heritability estimation (see for example, Elliott et al. 2018, Zhao et al. 2018). Here we replicate and follow up on our original results using a sample ten times larger, N = 26,818, which combines data from the UK Biobank project as well as 5 other projects that had previously participated in the ENIGMA consortium (IMAGEN, SHIP, TREND, Lothian, ADNI).

We used the same regional brain volumes estimates than those used in the ENIGMA consortium studies by Stein et al. (2012) and Hibar et al. (2015): several subcortical structures, total brain volume and intracranial volume. In addition, we also studied height and intelligence scores. Brain volume correlates with height and intelligence scores (Taki et al. 2012), which are both known to be heritable (Yang et al. 2010, Plomin and Deary 2015). We aimed thus at determining to which extent the heritability of regional brain volumes was given by their relationship with height (i.e., affected by the same genetic factors that determine body size), or if different genetic factors affected them specifically. Similarly for intelligence scores, we aimed at better understanding its relationship with brain volume.

For all phenotypes, we estimated to what extent genome-wide SNPs were able to capture the inter-individual variability in regional brain volumes, that is, we estimated the proportion of phenotypic variance captured by SNPs across the genome (also called SNP heritability). Additionally, we used GWAS data to compute genome-wide polygenic scores, which provide a phenotypic prediction at the individual level. The analyses of the influence of the complete genome on our phenotypes were complemented with analyses on a series of genomic partitions: genic versus non-genic; preferential expression in the central nervous system or by cell type; low, medium or high minor allele frequency. This type of analysis can reveal whether specific genomic regions are enriched in the amount of variance they capture. Finally, we looked at the pleiotropy across phenotypes. For this, we computed genetic correlations and phenotypic correlations for all pairs of phenotypes. For brain regions, we also compared the genetic and phenotypic correlations between the left and right parts of the same structure as a means to estimate the role of genetics and environment in brain asymmetry.

Our results confirm the observation that polygenic factors play an important role in the determination of neuroanatomical variability, and show different ways in which biological information can be obtained to better understand polygenic effects.

## Material and Methods

### Data sharing

We obtained whole-genome genotyping from N = 26,818 subjects from 6 different projects: UK Biobank, IMAGEN, ADNI, Lothian Birth Cohort 1936, SHIP and TREND. Extensive efforts have been made to homogenise the neuroanatomical measurements across sites, which were described in the ENIGMA 1 and 2 reports (Stein et al. 2012, Hibar et al. 2015), and the UK Biobank neuroimaging analysis group (Alfaro-Almagro et al. 2018). The UK Biobank project (https://imaging.ukbiobank.ac.uk) is a large, long-term biobank study in the United Kingdom aiming at investigating the contributions of genetic predisposition and environmental exposure to the development of disease. The study is following about 500,000 volunteers enrolled at ages from 40 to 69 years old, 54% females. IMAGEN (https://imagen-europe.com) is a project to identify and characterise specific genetically-influenced alterations in reinforcer sensitivity and executive control which manifest in adolescence and carry the risk for overt psychopathology later in life. It includes general population 13 to 17 year old adolescents (49% of females) from Germany, France, Ireland, and the United Kingdom. ADNI, the Alzheimer’s Disease Neuroimaging Initiative (http://adni.loni.usc.edu), is a longitudinal multicenter study designed to develop clinical, imaging, genetic, and biochemical biomarkers for the early detection and tracking of Alzheimer’s disease in the United States of America. The dataset combines data from the initial five-year study (ADNI-1), and the follow-ups ADNI-GO, ADNI-2, and ADNI-3. It includes subjects 54 to 90 years old, 42% female. The Lothian Birth Cohort 1936 (https://www.lothianbirthcohort.ed.ac.uk) is a follow-up of the Scottish Mental Surveys of 1947, which tested the intelligence of almost every child born in 1936 and attending school in Scotland in the month of June 1947. It includes subjects 71 to 73 years old, 47% female. The SHIP and TREND cohorts contain data from the Study of Health in Pomerania (SHIP, http://www2.medizin.uni-greifswald.de/cm/fv/ship.html), a population-based epidemiological study consisting of two independent cohorts SHIP and SHIP-TREND. These projects investigate common risk factors, subclinical disorders and diseases in a population of northeast Germany. The dataset included data from subjects 21 to 90 years old, 44% female for TREND, 48% female for SHIP. All data sharing was approved by our local ethical board as well as by those of the participating projects wherever required. The list of projects and their respective number of subjects is described in Table 1.

**Table 1.** Sample sizes and number of variants per project. *The UK Biobank dataset was split in two parts: N = 14,144 subjects were used for heritability analyses (13,086 included), and N = 6,678 subjects were used for validation of genome-wide polygenic scores (6,184 included).

| Project | Total sample size | Percentage of females | Mean age (standard deviation) | Sample size included | Total number of variants | Number of variants included |
| --- | --- | --- | --- | --- | --- | --- |
| IMAGEN | 2,011 | 49% | 14.6 (0.4) | 1,736 | 573,299 | 267,151 |
| Lothian Birth Cohort 1936 | 1,005 | 47% | 72.7 (0.7) | 544 | 529,015 | 256,417 |
| TREND | 858 | 44% | 50.0 (13.5) | 813 | 2,389,858 | 597,902 |
| SHIP | 963 | 48% | 56.5 (12.6) | 941 | 863,230 | 271,635 |
| ADNI | 1,189 | 42% | 74.2 (7.1) | 986 | 331,088 | 227,005 |
| UK Biobank* | 20,792 | 54% | 62.6 (7.5) | 19,270 | 734,447 | 490,061 |

### Regional brain volumes

The measurements of regional brain volume coming from projects that had previously participated in ENIGMA (IMAGEN, Lothian Birth Cohort 1936, TREND, SHIP and ADNI) were the same that had been used in Stein et al. (2012) and Hibar et al. (2015). For the UK Biobank subjects, the estimation of the volumes was performed using FreeSurfer 6.0 (https://surfer.nmr.mgh.harvard.edu). For comparison, we also included the estimates obtained using FSL FIRST (https://fsl.fmrib.ox.ac.uk/fsl) that were made available by UK Biobank (the processing pipeline is described in https://biobank.ctsu.ox.ac.uk/crystal/docs/brain_mri.pdf). In addition to the subjects excluded in the quality control made by the UK Biobank, we excluded 52 additional subjects showing an extreme relationship between total brain volume and intracranial volume. For this, we used a kernel density estimator to fit a probability density function to the intracranial versus brain volume data, and tagged as outliers all subjects with a local density inferior to 1% of the maximum density for UK Biobank, and 2% for other datasets.

The regions included in our analyses were: nucleus accumbens (labelled as Acc), amygdala (Amy), putamen (Pu), pallidum (Pa), caudate nucleus (Ca), hippocampus (Hip) and thalamus (Th); along with brain volume (BV) and intracranial volume (ICV). In addition to these regions, we investigated height and intelligence scores (IS), available from the UK Biobank and IMAGEN projects. It is important to note that the fluid intelligence score in UK Biobank (a 2 minute test aiming at evaluating the capacity to solve problems that require logic and reasoning ability, independent of acquired knowledge; see https://biobank.ctsu.ox.ac.uk/crystal/label.cgi?id=100027), is not the same as the intelligence score used by the IMAGEN project, which was obtained using the WISC test.

### Genotype filtering

All genetic analyses were performed for each project independently. Genotyping data was converted to the hg19 reference wherever required using UCSC LiftOver (http://genome.ucsc.edu/cgi-bin/hgLiftOver). We used genotyped autosomal SNPs (single nucleotide polymorphisms) which passed UK Biobank quality control for all batches (http://www.ukbiobank.ac.uk/wp-content/uploads/2014/04/UKBiobank_genotyping_QC_documentation-web.pdf). Additionally, we removed SNPs in 24 regions with long range linkage disequilibrium (LD, see Price et al. 2008). SNPs were then filtered to exclude those with minor allele frequency (MAF) <0.1%, missing rate >1%, or Hardy-Weinberg disequilibrium with a p<10^−6^. Individuals were removed when >10% of their SNPs were missing. We finally pruned SNPs which were in linkage disequilibrium with a variance inflation factor >10, which corresponds to a multiple *R*^2^ for the regression over linked SNPs <0.9. The filtering was made using PLINK v1.90b3.46 (Purcell et al. 2007).

### Genetic relationship matrices (GRM)

GRMs were computed based on autosomal chromosomes using GCTA v1.91.3 (Yang, Lee, et al. 2011). We included only one of each pair of subjects with an estimated relatedness >0.025 (approximately corresponding to cousins two to three times removed). GRMs were computed per chromosome and then merged for the whole genome.

### Population structure

Genetic variance estimates based on genomic estimates of relatedness are sensitive to cryptic relatedness and population structure. These factors can influence the phenotypic similarity beyond the estimated degree of genetic relatedness (Yang, Manolio, et al. 2011, Browning and Browning 2011). In addition to the exclusion of subjects with a degree of genetic relatedness greater than 0.025, we used the first 10 principal components of the GRM as covariates in our statistical analyses.

### Genetic variance

We estimated the amount of phenotypic variance captured by SNPs using a linear mixed model with age, sex, imaging centre, and the first 10 principal components of the GRM as fixed effect covariates, and a random effect with a covariance matrix corresponding to the GRM (GCTA GREML method, Yang et al. 2011). We estimated SNP heritability as the ratio of the genetic variance to the phenotypic variance, with genetic variance being the variance of the random component, and the phenotypic variance being the sum of random component and residual component with fixed effects removed. We used GCTA v1.91.3 (Yang, Lee, et al. 2011) for those computations and did not constrain genetic variance estimates to lie in the range 0% to 100%, in order to obtain unbiased estimates (option *--reml-no-constrain*).

### Genetic correlation

Genetic correlation was estimated using GCTA REML bivariate analysis (Lee, Yang, et al. 2012) in constrained mode (option *--reml-bivar*). Both phenotypic and genetic correlation were adjusted for age, sex, imaging centre, and the first 10 principal components of the GRM. We compared genetic and phenotypic correlations using the delta method to estimate standard errors (R package msm https://cran.r-project.org/web/packages/msm/). We report estimates with their standard errors (s.e.).

### Genetic variance partitioning

In its simplest form, GCTA allows to estimate the amount of variance captured by the matrix of genetic relationships, assuming that each SNP captures the same amount of variance. Through genomic partitions we can create different genetic relationship matrices based on non-overlapping regions of the genome. The SNPs on each of these partitions can capture then a different amount of variance (although, as before, SNPs within a given partition are supposed to capture all the same amount of variance). We grouped SNPs in the following partitions:

1. Partition based on genic status. Using 66,632 gene boundaries from the UCSC Genome Browser hg19 assembly, we made a first set with all SNPs within these boundaries, two further sets that included also SNPs 0 to 20 kbp and 20kbp to 50 kbp upstream and downstream of each gene, and a last set including the SNPs not located in regions less than 50 kbp upstream or downstream of genes. Both exonic and intronic regions were included in the genic regions. These partitions do not correspond exactly to those used by Toro et al. (2015) which were: one with strict genic/non-genic boundaries (0 kbp), another with genic ± 20 kbp versus the rest, and finally genic ± 50 kbp versus the rest.
2. Partition based on preferential central nervous system (CNS) expression (Raychaudhuri et al. 2010, Lee, DeCandia, et al. 2012) using ± 50 kbp as gene boundaries, and based on markers of brain cell types as defined by two recent scRNA-Seq studies (Skene et al. 2018, 2018, Li et al. 2018) using ± 20 kbp as gene boundaries.
3. Partition based on allele frequency. A partition based on MAF with 4 groups: from 0.1 to 5%, from 5 to 20%, from 20 to 35% and from 35 to 50%. In Toro et al (2015), only the last 3 partitions were included, covering the range from 5 to 50% of MAF.

### Genetic variance partition enrichment

Once the variance captured by each partition was computed, we were able to estimate the significance of the difference in variance captured by an individual partition against a model where each SNP captures exactly the same amount of variance (a null hypothesis of no enrichment). In this latter case, the amount of variance captured by each partition should be directly proportional to the number of SNPs it contains.

We tested whether any of the partitions captured more variance than what could be expected given its number of SNPs. A Z-score was computed by comparing the SNP-set genetic estimated variance 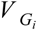 of partition *i* to the SNP-set genetic variance 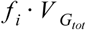 expected under no enrichment:

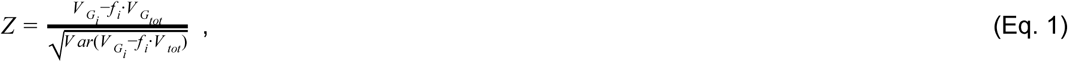

where *f* _*i*_ is the fraction of the SNPs included in partition *i*. We estimated the variance of the observed enrichment as:

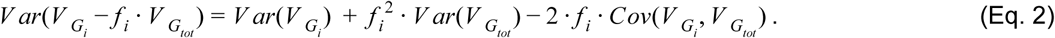

Here, 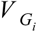 represents the genetic variance of partition *i* and 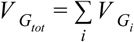. We preferred the analytic estimation over the estimation by permutations, because the permutation methods presented some limitations due to the difficulty of preserving the SNP structure of the partitions and due to the computational resources needed for the permutations (Fig. S1, S2).

### Meta-analytical combination of the estimates of each project

The independent estimates obtained from each of the projects were combined into a single one using an inverse variance weighting method. We validated the meta-analytical approach by comparing the distribution of genetic variance estimates from simulated heritable phenotypes with their theoretical normal distribution in sub-samples of ADNI and UK Biobank datasets.

### Accounting for measurement error

The estimation of regional brain volumes, height and intelligence scores is submitted to measurement errors. In particular, small, poorly defined brain regions are more difficult to measure accurately than larger ones. Measurement errors enter as environmental variance in the decomposition of phenotypic variance into genetic and environmental variance, and hence decrease the heritability estimates (*V* _*G*_ /*V* _*P*_). Heritability estimates unbiased for measurement errors can be obtained with repeated measurements (Ponzi et al. 2018), however, the datasets we used to measure brain volumes provide only one MRI per subject. We used datasets from CoRR, the Consortium for Reliability and Reproducibility (Zuo et al. 2014), which includes multiple MRIs per individual to estimate intraclass correlation coefficients (ICC) (Fisher 1970) for each regional volume. We segmented the volumes as we did in the UK Biobank dataset, using FreeSurfer 6.0. ICCs were computed on the volumes of 836 subjects with two MRI sessions after covarying for age, sex and scanning site. For intelligence, we estimated ICC from 1,301 subjects in UK Biobank who took the fluid intelligence test 3 times. The ICC for height was considered to be equal to 1 (no measurement error). Adjusted phenotypic variance and covariance estimates were obtained by multiplying the raw estimates by their corresponding ICC (see Supplemental Materials).

### Genome-wide polygenic scores

We used the SNP effects estimated in the association analysis of 13,086 UK Biobank subjects to estimate the phenotypes of 6,184 additional unrelated subjects with MRI data from the latest release of the UK Biobank project. The scores were estimated from the filtered SNPs (not LD pruned). SNPs under various association p-value thresholds were selected and the ones in LD with a more significantly associated SNP were clumped. The p-value threshold that produced the best fit with the target dataset was selected. We used the software PRSice associated with PLINK for the computation of genome-wide polygenic scores (Euesden, Lewis, and O’Reilly 2015). Each phenotype was regressed on age, sex, scanning centre, and the 10 first principal components of the GRM. The analyses were performed on the residuals of this linear regression. We then estimated the proportion of variance captured by genome-wide polygenic scores in each phenotype using the coefficient of determination *R*^2^. For comparison, we also computed the predicted height of the 6,184 subjects using SNP effects from an independent group of 277,756 unrelated (with a pairwise estimated relatedness <0.025 in the GRM) UK Biobank subjects.

## Results

### Genome-wide variants capture a large proportion of the diversity of regional brain volumes, height and intelligence scores

The heritability estimates for intracranial volume, total brain volume, as well as the volume of subcortical structures were substantial and statistically significant (Fig. 1, Table S1.1). Estimates for intracranial volume and total brain volume were large: V_G_/V_P_(ICV) = 53 ± 4.5% (all our variance estimates are reported as estimation ± s.e.), V_G_/V_P_ (BV) = 52 ± 4.5%. Similarly, the genetic variance estimates for subcortical structures were all above 40%: accumbens (V_G_/V_P_ = 40 ± 4.5%), amygdala (V_G_/V_P_ = 45 ± 4.5%), putamen (V_G_/V_P_ = 48 ± 4.5%), palladium (V_G_/V_P_ = 40 ± 4.5%), caudate (V_G_/V_P_ = 52 ± 4.5%), thalamus (V_G_/V_P_ = 54 ± 4.5%), hippocampus (V_G_/V_P_ = 53 ± 4.5%). All estimates were highly statistically significant with p<10^−11^ in all cases (log-likelihood ratio statistics from 49 to 242 in UK Biobank alone) (Fig. 1, Table S1.2). The V_G_/V_P_ estimate for caudate nucleus (V_G_/V_P_ = 52 ± 4.4%) was statistically significant and very different from what was observed in the IMAGEN cohort (V_G_/V_P_ = 4 ± 24%) which, as mentioned in Toro et al. (2015), may reflect an age bias specific to IMAGEN (where individuals were on average 14 years old).

**Figure 1.**
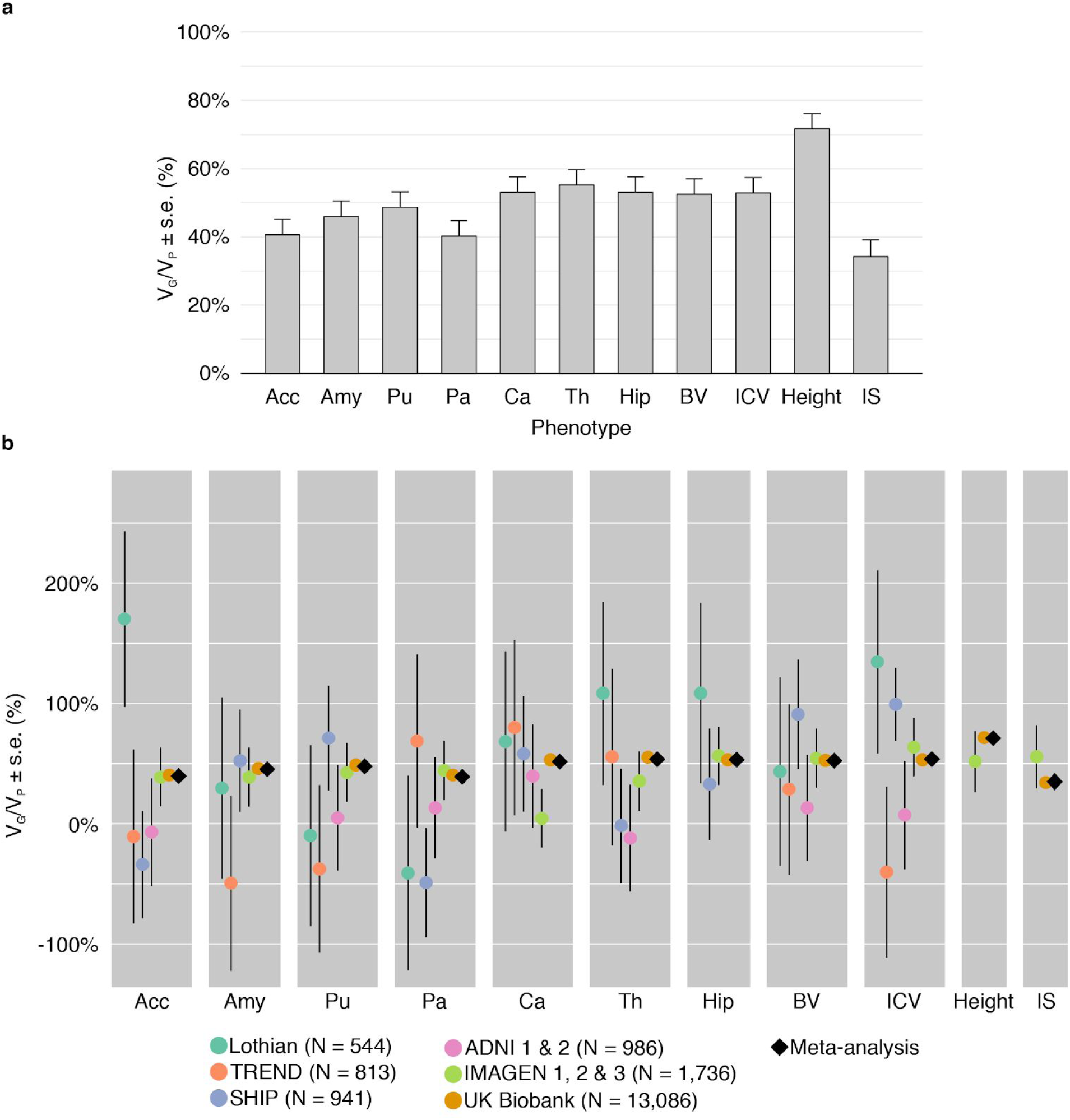
**a.** Proportion of variance captured by common genotyped variants (V_G_/V_P_) for brain regions, height and intelligence score. Meta-analytic estimates were obtained using inverse variance weighting of the estimates of the different projects studied. **b.** V_G_/V_P_ estimates for each project. The estimates were obtained using GCTA, without constraining the results to lie in the 0% to 100% range. The diamond shows the meta-analytic estimation. Age, sex, center, and the first 10 principal components were included as covariates. The error bars show the standard errors of the V_G_/V_P_ estimates.

The heritability estimate (V_G_/V_P_) for height (which combines only UK Biobank and IMAGEN) was large: V_G_/V_P_ = 71 ± 3.1% and close to those obtained from twin studies (Polderman et al. 2015) and from previous SNP heritability estimates (V_G_/V_P_ = 68.5 ± 0.4% in Ge et al. 2017). The fluid intelligence score in UK Biobank had the lowest heritability among all 11 studied phenotypes, V_G_/V_P_ = 35 ± 4.8%. This estimate is similar to the one obtained by Davies et al. (2011) on 30,801 UK Biobank subjects (V_G_/V_P_ = 31% ± 1.8%). The verbal and performance intelligence scores (VIQ and PIQ) in IMAGEN, although not based on the same test as fluid intelligence (FI) in UK Biobank, aim at capturing a similar phenotype. The estimate for fluid intelligence in UK Biobank seem smaller than those for verbal and performance intelligence quotients in IMAGEN (∼56 ± 26%), however, they were not statistically significantly different.

Our estimates were obtained using a meta-analytic approach to combine the results of the different projects using inverse variance weighting. In all cases, the meta-analytic estimates closely corresponded with the values obtained for the UK Biobank project (Table S1.2), because of its large sample size which accounted for ∼94% of the weighted estimates. Standard errors agreed well with the theoretical values proposed by Visscher et al. (2014), and implemented in the GREML statistical power calculator (http://cnsgenomics.com/shiny/gctaPower): they were from 1.01 to 1.05 times higher than the theoretical values when taking into account the variance of the genetic relationships, except for ADNI where they were 1.21 times higher). As expected, the standard errors decreased with sample size, from about 76% for ∼550 individuals, 43% for ∼1,000 individuals, 24% for ∼1,750 individuals (IMAGEN), down to 4% for UK Biobank with more than 13,000 individuals. Our result from simulated phenotypes showed that the estimates of V_G_/V_P_ obtained with sub-samples of N = 800, N = 400, N = 200 and even N = 100 subjects from the UK Biobank project were unbiased relative to the expected normal distribution. The simulations based on the ADNI project, however, showed a significant bias towards positive values when the sub-samples included N = 100 to N = 400 subjects, probably due to the heterogeneity of the population (Figures S12-13 and Table S6-7).

Similarly to what we had observed previously (Toro et al. 2015), the genetic variance estimates were not significantly affected by population structure: the non-inclusion of the 10 first PCs did not impact the estimates of variance, which changed on average by less than 4% (Fig. S3).

Finally, our heritability estimates for the UK Biobank and ADNI projects were comparable to those of previous studies of these projects by Elliott et al. (2018) and Zhao et al. (2018) (see Fig. S6 and Supplemental Materials).

### Genetic partitions show significant enrichment of the proportion of variance captured by specific gene sets

#### Partition per chromosome (autosomes)

We observed a strong correlation between V_G_/V_P_ estimates and chromosome size, which was significant for all phenotypes except the pallidum (Fig. 2). The correlation coefficients ranged from 0.30 ± 0.21 to 0.80 ± 0.13, r = 0.63 on average, capturing 41% of the variance (estimated as *R*^2^).

**Figure 2.**
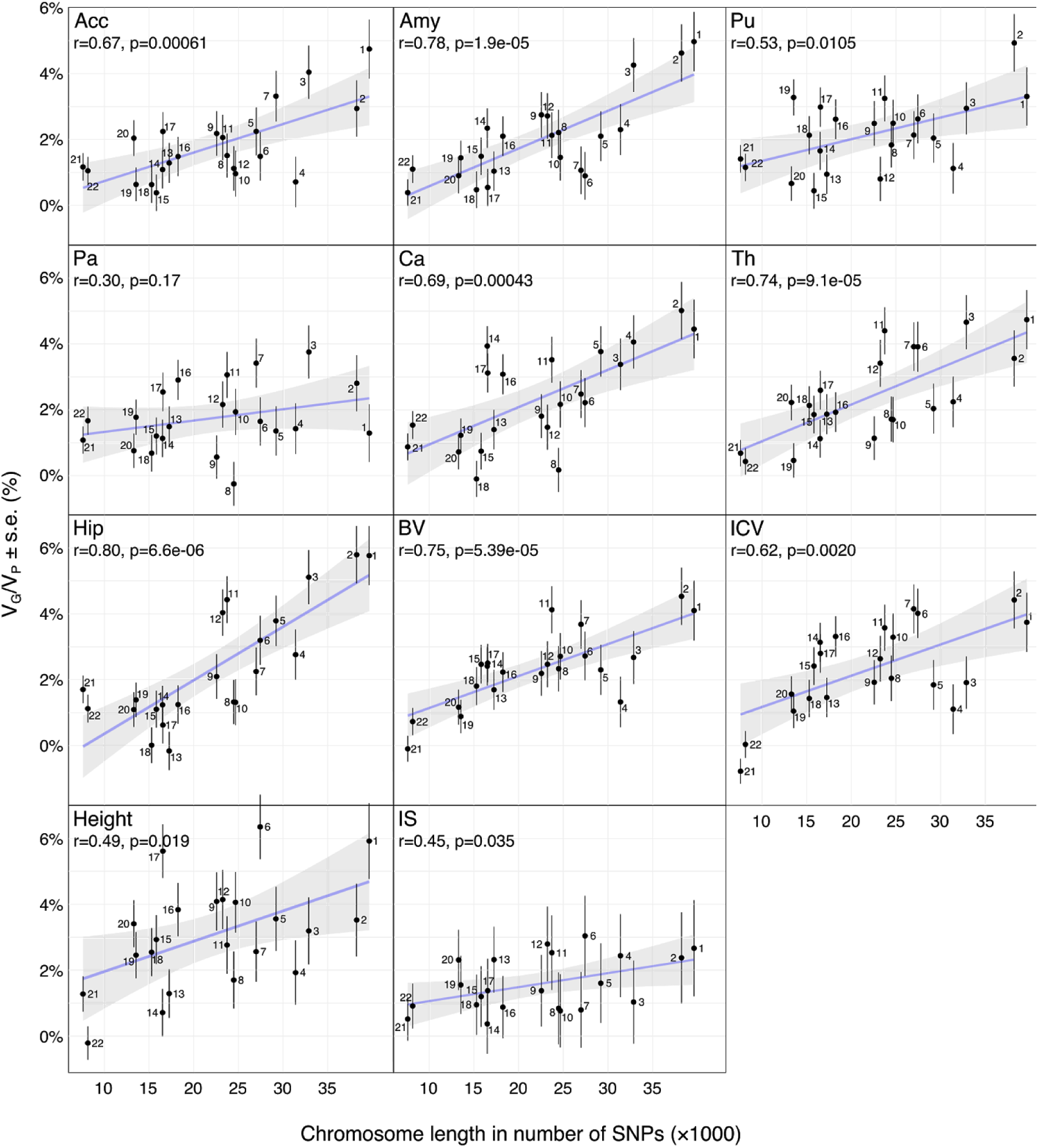
Scatter plots of the number of SNPs per chromosome versus V_G_/V_P_ estimates computed for each chromosome. V_G_/V_P_ estimates were obtained by partitioning SNPs across chromosomes and computed using the GCTA REML unconstrained method for total subcortical volumes. Age, sex, center, and the top 10 principal components were included as covariates. The error bars show the standard errors of the V_G_/V_P_ estimates.

#### Partition between genic and non-genic regions

The genic SNP set (± 0 kbp) contained 51% of all genotyped SNPs and captured on average 69% of the variance attributable to SNPs of most of the studied phenotypes, significantly more than what we would expect from its length (FDR<5%). The only exceptions were fluid intelligence and nucleus accumbens, for which it explained respectively 59 ± 9% and 55 ± 6% of the total genetic variance (Fig. 3, Table S4). Height was the only phenotype for which we observed an enrichment of V_G_/V_P_ captured by one of the non-genic SNP sets: the set of SNPs within 0 ± 20 kbp of the genic set. This set contained 15% of all SNPs but explained 27% of the variance of the height phenotype attributable to SNPs (FDR corrected p<0.01). In total, the variants located between 0 and 50 kbp away from genes captured 36 ± 4% of the genetic variance of height (FDR corrected p<0.05).

**Figure 3.**
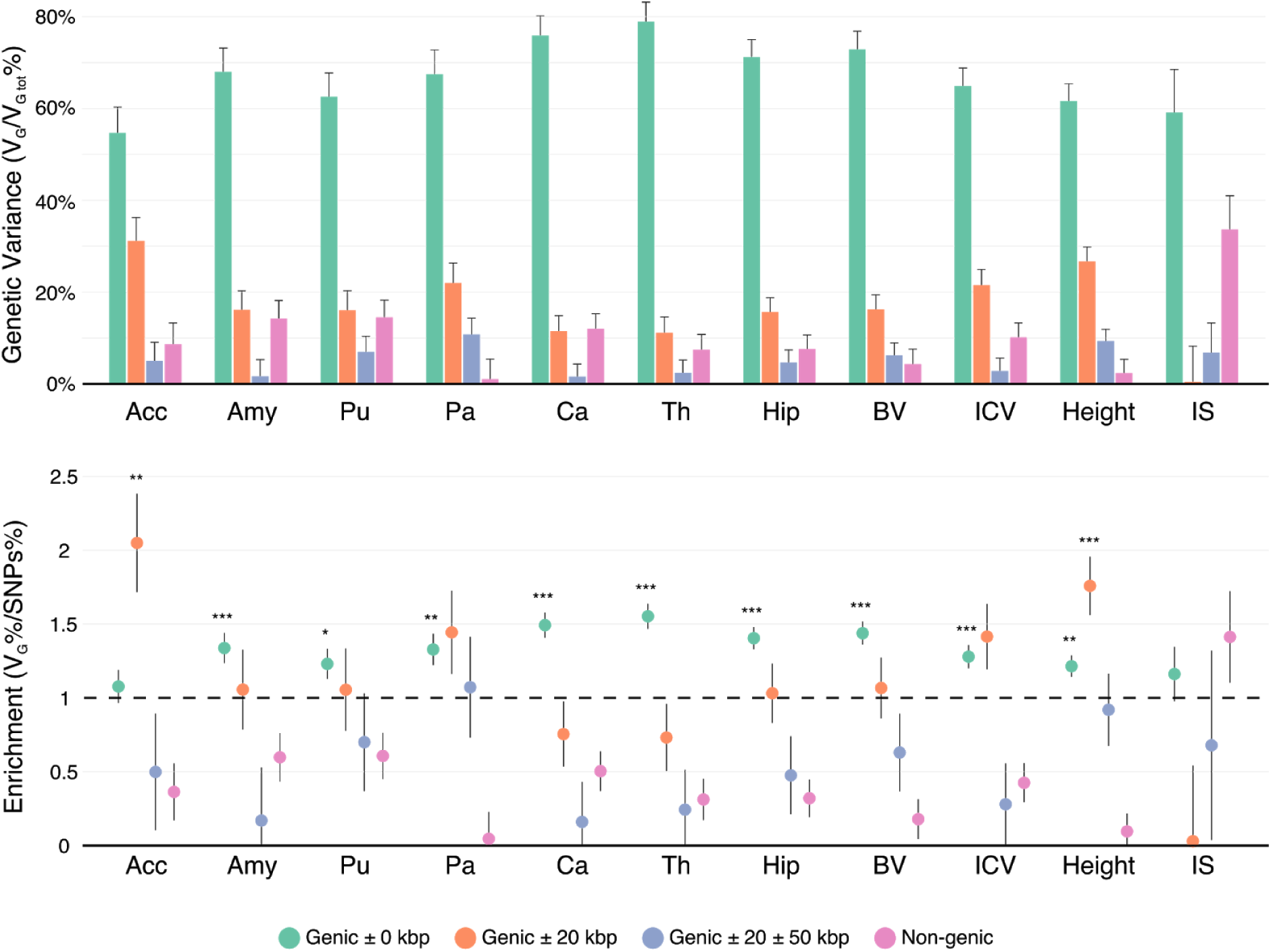
Variance enrichment for partitions based on closeness to genic regions. Meta-analytic estimates were obtained using inverse variance weighting of the estimates of the different projects studied. Top: V_G_/V_Gtot_ estimates computed for four sets of SNPs based on their distance to gene boundaries: all SNPs within the boundaries of the 66,632 gene boundaries from the UCSC Genome Browser hg19 assembly, two further sets that included also SNPs 0 to 20 kbp and 20kbp to 50 kbp upstream and downstream of each gene, and a remaining set containing SNPs located farther than 50kb from one of the gene boundaries. V_G_/V_Gtot_ estimates were computed using the GCTA REML unconstrained method for height, intelligence, and brain, intracranial and total subcortical volumes. The error bars represent the standard errors. Bottom: Enrichment of variance captured by each partition. The y-axis shows the ratio of the fraction of genetic variance explained by each partition divided by the fraction of SNPs contained in each partition. If all SNPs explained a similar amount of variance, this ratio should be close to 1 (dashed line). A Z-test was used to compare the ratios to 1 and p-values were FDR adjusted (\**p*<0.05, \*\**p*<0.01, \*\*\**p*<0.001).

#### Partition by involvement in preferential CNS expression and in brain cell types

No statistically significant enrichment was observed for CNS expression nor brain cell type partitions.

#### Partition by allele frequency

SNPs within the low MAF partition (MAF<5%) captured less genetic variance per SNP than those with medium and high MAF (the 3 partitions with MAF>5%) (Fig. 3), as previously described by Speed et al. (2017). Our four MAF-based partitions included the following average proportions of total SNPs: (1) MAF from 0.1% to 5%: comprising 40% of SNPs, (2) 5% to 20%: with 30% of SNPs, (3) 20% to 35%: with 12% of SNPs, and (4) 35% to 50%: with 9% of SNPs. Although the partition of SNPs with low MAF contained 40% of the SNPs, it captured on average only about 16% of the total genetic variance. This is less than expected in the GCTA model where each SNP captures the same amount of phenotypic variance, but slightly more than expected in a neutral theory of evolution, where the captured variance is proportional to the size of the MAF bin (Yang et al. 2017, Visscher et al. 2012).

### Genetic factors influencing volume are shared among brain regions

We observed extensive pleiotropy across brain regions, with an average genetic correlation of r_G_∼0.45 (Fig. 4). Genetic correlations, which represent the correlation between genetic effects of two phenotypes, were computed for each pair of brain regions, ICV, Height and IS, using data from the UK Biobank project. Figure 4 shows the genetic correlation matrix together with the phenotypic correlation matrix (see also Table S5).

**Figure 4.**
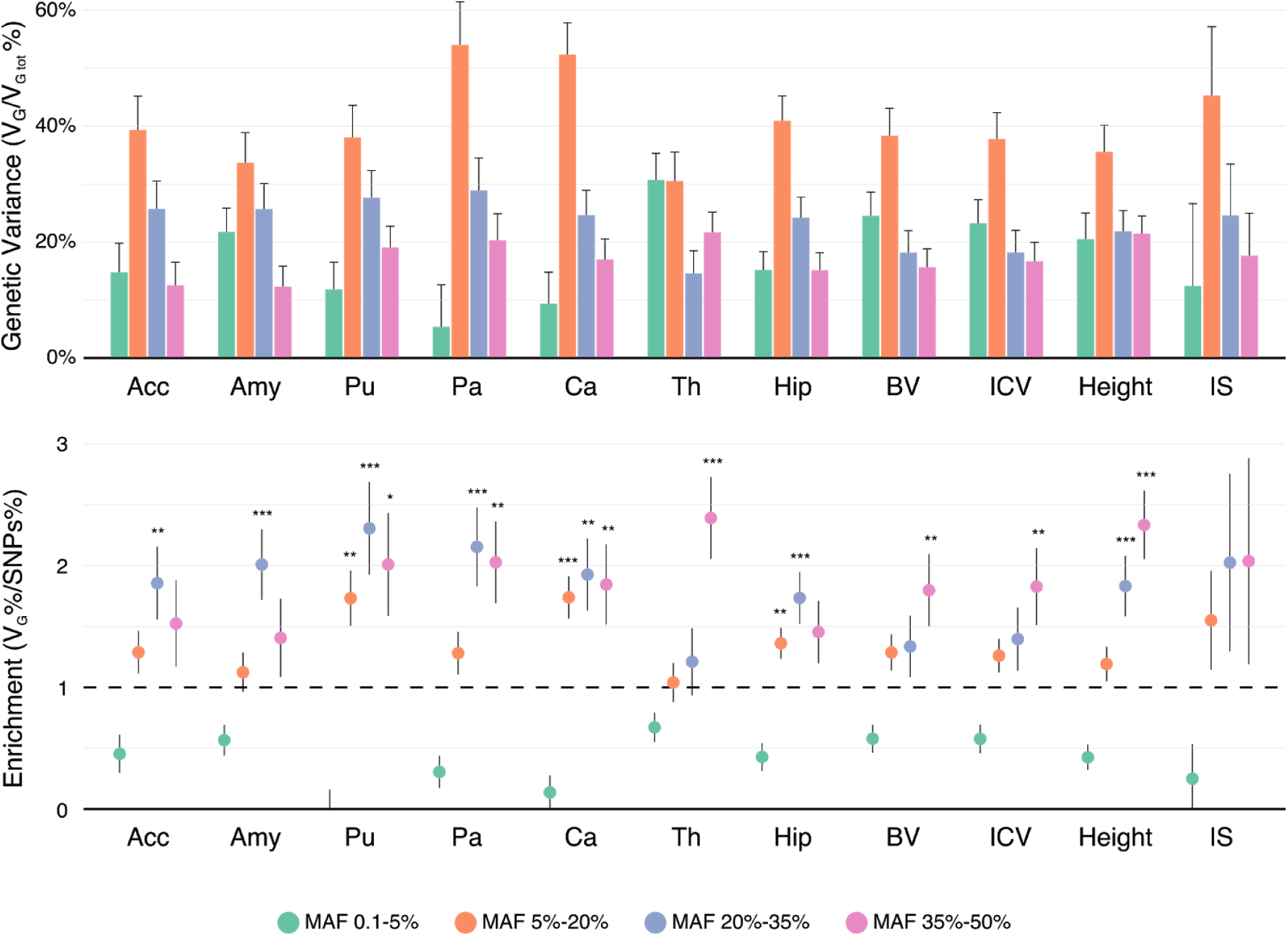
Variance enrichment for partitions based on minor allele frequency (MAF). Meta-analytic estimates were obtained using inverse variance weighting of the estimates of the different projects studied. Top: V_G_/V_Gtot_ estimates computed for four sets of SNPs based on their MAF: from 0.1 to 5%, from 5 to 20%, from 20 to 35% and from 35 to 50%. V_G_/V_G tot_ estimates were computed using the GCTA REML unconstrained method for height, intelligence, and brain, intracranial and total subcortical volumes. The error bars represent the standard errors. Bottom: Enrichment of variance captured by each partition. The y-axis shows the ratio of the fraction of genetic variance explained by each partition divided by the fraction of SNPs contained in each partition. If all SNPs explained a similar amount of variance, this ratio should be close to 1 (dashed line). A Z-test was used to compare the ratios to 1 and p-values were FDR adjusted (\**p*<0.05, \*\**p*<0.01, \*\*\**p*<0.001).

Genetic correlations were in general similar to phenotypic and environmental correlations, however, genetic correlations were often larger than phenotypic correlations for the left/right volumes of the same region. The concordance between phenotypic and genetic correlation was high (*R*^2^ = 0.80, Fig 5c), consistent with the report by Sodini et al. (2018) for other traits, and the correlation matrices were similar (Fig. S7). Phenotypic correlations can be decomposed as a sum of genetic and environmental correlations. The concordance between genetic and environmental correlations was also strong (*R*^2^ = 0.47, Fig. S8).

**Figure 5.**
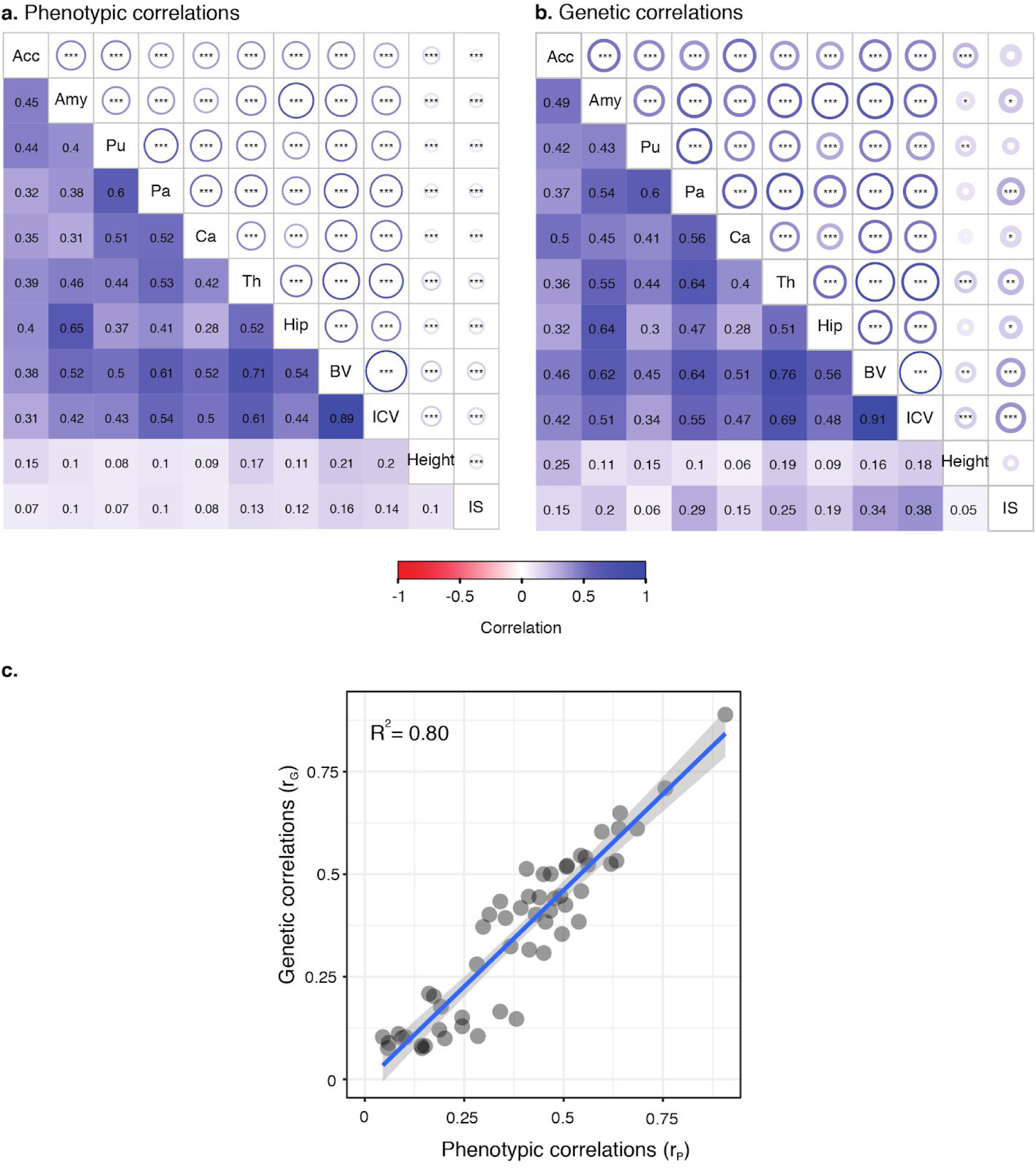
Phenotypic and genetic correlations. Significant phenotypic (a) and genetic (b) correlations were observed for most phenotypes. Correlation estimates are shown in the lower triangular part of the matrices, statistical significance in the upper triangular part. Circle radius represents correlation strength, stars indicate statistical significance of the correlation being non null (\**P*<0.05, \*\**P*<0.01, \*\*\**P*<0.001). The scatter plot (c) of phenotypic versus genetic correlations.

### Environmental factors are important in shaping brain asymmetry

For all brain regions, with the exception of the nucleus accumbens, the differences between their left and right volumes appeared to be of environmental origin. When considering total volumes (left plus right), the differences between genetic and phenotypic correlations were not different from zero (Z-test FDR > 50%, Table S5). However, the situation was different when considering the regional volume asymmetries (left minus right). In that case, the genetic correlations of the asymmetries were not statistically different from r_G_ = 1, and were in all cases statistically significantly larger than the phenotypic correlations (Z-test FDR<5%, Fig. S9 and Table S5), which suggests an important role of the environment in shaping hemispheric asymmetry. To further test this hypothesis, we measured the V_G_/V_P_ of the differences between left and right volumes of each structure. Only the heritability of the volume asymmetry of the nucleus accumbens was significantly different from zero (V_G_/V_P_ = 16 ± 4.5%). Except for this structure, the differences in volumes between right and left hemispheres of all the other brain regions seemed to be only of environmental origin.

### Genetic factors influencing brain volume are shared with height and intelligence scores, but less between height and intelligence scores

The genetic correlation between height and brain volume (r_G:BV.height_ = 0.164 ± 0.053) was smaller than the genetic correlation between brain volume and intelligence (r_G:BV.IS_ = 0.343 ± 0.080). However, the difference was not statistically significant: r_G:BV.IS_ - r_G:BV.height_ = 0.179 ± 0.095, 95% CI: −0.008, 0.365 (computed using the formula xxxvii in Pearson and Filon 1898). The phenotypic and genetic correlations between intelligence scores and height were, however, the smallest we observed across all phenotypes: r_P_ = 0.101 ± 0.009 and r_G_ = 0.048 ± 0.073. This is different to what we had previously observed in the IMAGEN cohort (Toro et al. 2015), where the correlation between brain volume and intelligence scores was higher than the correlation between brain volume and height.

### Heritability and genetic correlation estimates remain statistically significant after accounting for measurement errors

The estimates of heritability, phenotypic and environmental correlation can be biased by noise in the volume measurements, which could in particular explain the differences between genetic and phenotypic correlations of left and right volumes. We investigated this hypothesis by removing the effects of measurement errors estimated from the Consortium for Reliability and Reproducibility (Zuo et al. 2014) for FreeSurfer segmentations, and from the repeated measures of intelligence score in UK Biobank. Intraclass correlation coefficient (ICC) for intelligence scores was estimated at 63% whereas the lowest ICCs for subcortical structures were found for accumbens (83%), pallidum (87%) and amygdala (89%). After correction for measurement error, height remained the most heritable phenotype although it was the only one that we did not correct. Disattenuated estimates of V_G_/V_P_ for intelligence became 54 ± 7.8%, which is similar to the disattenuated heritabilities of brain volumes (between 46 ± 5.3% for pallidum and 61 ± 5.0% for thalamus) (Fig. S10). Genetic correlations between left and right hemispheres of subcortical structures remained statistically significantly greater than the adjusted phenotypic correlations except for pallidum and accumbens (Table S5 and Fig. S11). Estimates of V_G_/V_P_ for the differences between left and right volumes remained low and not significantly different from 0 for all subcortical structures (between 5.4 ± 14% for pallidum and 16 ± 11% for thalamus) except for accumbens (34 ± 9.8%).

### Genome-wide polygenic scores captured a statistically significant but small proportion of phenotypic variance

Genome-wide polygenic scores based on a GWAS of 13,086 subjects captured a statistically significant although very small proportion of brain region volume variance. The genome-wide polygenic scores were computed for ∼6,000 additional participants from the UK Biobank who were not used in the GWAS. The predictions captured an amount of phenotypic variance ranging from 0.5% (p < 10^−22^) for the amygdala to 2.2% (p < 10^−31^) for brain volume (Fig. S15, Fig. S16, and Supplemental Table S8). For height, a genome-wide polygenic score obtained based on GWAS summary statistics from the 13,086 UK Biobank subjects with MRI captured ∼3% of the variance. To evaluate the impact of the number of samples used in the GWAS on the amount of variance captured by the genome-wide polygenic scores, we also computed the genome-wide polygenic scores for height using the GWAS summary statistics from ∼277k unrelated UK Biobank subjects not included in the validation dataset. This allowed us to capture ∼27% of the variance of height (an >8-times increase) (Fig. S17).

## Discussion

Our results suggest that neuroanatomical diversity is the product of a highly polygenic architecture, with SNPs capturing from 40% to 54% of regional brain volume variance, confirming our original findings (Toro et al., 2015) as well as those of others (Elliott et al. 2018, Zhao et al. 2018). At a global scale, causal variants were distributed across the genome: for different brain regions, chromosomes containing a larger number of SNPs captured a proportionally larger amount of variance than smaller chromosomes, with a correlation of r∼0.64 on average. At a local scale, however, SNPs within genes (∼51%) captured ∼1.5 times more genetic variance than the rest; and SNPs with low minor allele frequency (MAF) captured significantly less variance than those with higher MAF.

When partitioning genetic variance into MAF bins, the lowest MAF partition, going from 0.1% to 5% (10% of the total MAF range) contained ∼40% of all SNPs, but captured only ∼16% of the total genetic variance: SNPs with low MAF captured significantly less variance than those with higher MAF. However, they captured more variance than expected under a neutral evolution model, where a MAF bin is expected to capture an amount of variance proportional to its size (10% size but 16% of the variance). This result suggests a negative selection model, where loci of large effect are being removed from the population. This apparent contradiction between high effect size and low captured variance for low MAF variants can be explained by the fact that the relationship between captured heritability *h*^2^ and the allele effect *b* for a given SNP is dependent of its allele frequency *p* : *h*^2^ = 2 *b*^2^ *p* (1 − *p*) (Schoech et al. 2017, Zeng et al. 2018). Given this heterogeneity in effect sizes, using more flexible models that do not make strong assumptions on the relationships between effect size, LD, and MAF, such as GREML-LDMS (Yang et al. 2015), should improve the accuracy of the heritability estimates. Our study of rare variants is limited because of the use of datasets based on SNP arrays. The availability of whole-genome sequencing data for large cohorts is starting to allow the study of more refined partitions of rare variants and is showing that rare causal variants might be a main source of the variance remaining to be explained (Wainschtein et al. 2019).

In addition to showing the heritability of regional volume diversity, our analyses also show an extensive pleiotropy across brain regions. The computation of genetic covariance for pairs of brain regions allowed us to estimate their genetic correlation. We observed an average genetic correlation of r_G_∼0.45. Interestingly, we observed that although genetic correlations were similar to phenotypic correlations across brain regions, if we compared the left and right aspects of the same brain region, their genetic correlations were close to 1 and systematically larger than phenotypic correlations. This could be an indication that the observed phenotypic asymmetries in regional brain volume are of environmental origin. To confirm this result, we used an alternative way of analysing the genetic/environmental nature of regional asymmetry, by looking at the heritability of the differences in left/right volumes. These heritabilities were generally not statistically significantly >0, which again supports the idea of the environmental nature of regional brain asymmetries.

The analysis of genetic correlations also allows us to explore the link between neuroanatomical diversity and other anatomical parametres or even cognitive functions. Brain volume is correlated with body size, and in the recent years the polygenic architecture of height has been well described (Yengo et al. 2018). It could be argued that neuroanatomical diversity is simply determined by the same genetic factors that produce body size variability in general. The genetic correlation between brain volume and height was, however, relatively small (r_G_∼0.16), suggesting little overlap between their genetic causes (in Toro et al. 2015 we did not find a significant genetic correlation, most likely due to lack of statistical power). By contrast, the genetic correlation between brain volume and intelligence scores was of medium strength (r_G_∼0.34), suggesting a larger overlap. The difference between these two genetic correlations was not significant and the question deserves further study. The genetic correlation between height and intelligence scores was the smallest of all those we studied, r_G_<0.05, suggesting that their relationship with brain volume may be due to different genetic factors.

One important source of bias in our estimates of heritability and genetic correlation could be related to measurement error. In MRI data, for example, some brain regions are more clearly delimited than others, which makes them easier to segment accurately. Furthermore, there is an important variability in volume across brain regions. Segmentation errors would then be comparatively more important for small, poorly delimited regions such as the amygdala, than for large regions like the thalamus. We sought to take into account errors in automatic segmentation of brain regions by analysing data from the CoRR project (Zuo et al. 2014), where the same subjects were scanned several times. For intelligence scores, we used the subset of UK Biobank where subjects passed the fluid intelligence test on multiple occasions. Our results remained for the most part unchanged after adjusting the estimates for phenotypic measurement errors. A limitation of this approach, however, is that we used MRI scans from a different project (although the processing pipeline was the same). Ideally, one would have preferred to have repeated scans for as subset of UK Biobank subjects, because segmentation quality depends on MRI quality which varies between datasets. The availability of repeated measurements in the same datasets from which genetic variance is estimated may allow to more precisely distinguish between the different sources of phenotypic variations.

Another possible source of bias could be the multi-centric nature of our data. The results presented here come from six different projects. Rather than combining our genetic raw data into a single dataset, we chose to estimate the part of phenotypic variance captured by SNPs independently in each dataset and then combine estimates in a meta-analysis. While our chosen solution helps to handle heterogeneity, it trades on statistical power. The reason for this is that the standard error of the GCTA GREML heritability estimate is approximately inversely proportional to the number of subjects (Visscher et al. 2014) (Fig. S14). In the present analysis, the UK Biobank project accounted for ∼94% of the estimates, largely driving the results. Indeed, the inclusion of the other projects reduced the standard error of the estimates based on the UK Biobank project only by a factor of ∼1.03. If the raw genotyping data had been combined in a single mega-analysis instead of in a meta-analysis, the decrease in standard error would have been ∼1.36 smaller than what we reported. The meta-analytical approach may however prove interesting in the future when large imaging genetics datasets other than UK Biobank will be available or in cases where raw genotyping data cannot be easily shared.

Our analyses are also limited in their ability to provide information at the individual level. With the advent of large GWAS studies, genome-wide polygenic scores (GPS) have become increasingly used to predict phenotypes from whole-genome genotyping data. GPSs are based on effect sizes estimated through GWAS in a large population. These estimates can then be used to predict the phenotype in an independent sample. We aimed at evaluating to which extent GWAS of the N∼13k UK Biobank subjects used for our heritability analyses allowed us to predict the phenotypes of additional N∼6k subjects. GPSs captured a very small, although statistically significant, proportion of the variance. For brain volume, for example, GPS captured ∼2.5% of the variance. This is the expected result in the presence of a strongly polygenic phenotype (Wray et al. 2013). It is also expected that prediction accuracy will improve as the number of subjects used for effect size estimation increases (the UK Biobank project alone should provide data for N∼100k in the years to come). We aimed at testing the potential increase in predictive power by computing a GPS for height. When effect sizes were estimated from N∼13k subjects, GPS captured ∼3% of height variance, however, when effect sizes were estimated from N∼277k subjects, the amount of variance explained increased to ∼27%. We expect that effect sizes for regional brain volume estimated from N∼100k will allow us to compute GPSs capturing ∼12% of the variance (based on Wray et al. 2013 and Daetwyler, Villanueva, and Woolliams 2008).

The detection of candidate genes of large effect is an appealing tool for gaining mechanistic insight on normal and pathological phenotypes. However, this approach is ill adapted to strongly polygenic architectures, where not only a few large-effect alleles are involved, but potentially hundreds of thousands of alleles of almost infinitesimal effect (Wray et al. 2018). Neuroimaging endophenotypes such as those obtained using structural and functional MRI could provide an alternative source of mechanistic insight. The brain imaging literature is rich in examples of associations between different brain regions and networks with normal and pathological cognitive phenotypes. The automatic mining of these associations could provide a layer of annotation for brain regions and networks similar to those available today for genome annotation. Further investigation of the genomic architecture of neuroimaging endophenotypes should prove an important tool to better understand the biological basis of brain diversity and evolution in humans, as well as the biological basis of the susceptibility to psychiatric disorders.

## Supporting information

h2 all datasets maf001

h2 all SNPs

h2 partitions Freesurfer

h2 partitions, merged

correlations

h2 simulations, adni

h2 simulations, UKB

Polygenic scores, UKB

