## Supplemental Material for "Polygenic architecture of human neuroanatomical diversity"

**Supplementary Material**

### 1. Comparison of methods for the estimation of enrichment in genetic variance partitioning.

We can estimate the significance of the difference in variance captured by an individual partition against a model where each SNP captures exactly the same amount of variance. In this latter case, the amount of variance captured by each partition should be directly proportional to the number of SNPs it contains.

We first aimed to assess the statistical significance of the partition enrichment using a permutation test as in Toro et al. (2015) and a derivation of statistical significance obtained from the covariance matrix of variance estimates reported by GCTA.

In the permutation test, SNPs were randomly selected to build the partitions, keeping the same number of SNPs as in the original partition. For each set of random partitions, the same linear mixed-effects model as before was fitted (including age, sex, centre and 10 PCs). We tested whether any of these partitions captured more variance than what could be expected given its number of SNPs. Briefly, a Z-score was computed by comparing the SNP-set genetic estimated variance $V_{G_i}$ of partition i to the SNP-set genetic variance $f_i \cdot V_{G_{tot}}$ expected under no enrichment:

$$Z = \frac{V_{G_i} - f_i \cdot V_{G_{tot}}}{\sqrt{Var(V_{G_i} - f_i \cdot V_{tot})}} \qquad \text{(Eq. 1)}$$

where $f_i$ is the fraction of the SNPs included in partition i. The p-value of enrichment was computed by comparing the observed Z score to those obtained from 1,000 permutations.

One limitation of this approach is that it does not preserve the local LD relationships among SNPs. We observed that the standard errors of the genetic variance estimates reported by the permutation approach were systematically larger than those computed from the covariance matrix of variance estimates. To better preserve the original LD, we tried an alternative permutation method in which we permuted blocks of contiguous SNPs. For the partitioning based on genic status, the standard errors of the simulated partitions estimates were compatible with the standard error of the original partitions (Fig. S1). However, this alternative permutation method did not reduce the gap with the theoretical values for the partitioning by MAF since they depend only on the frequency of the individual SNPs, and not on their contiguity over the genome (Fig. S2). Because of this, and because the permutation approach was much more demanding in terms of computation, we decided to use only the theoretical derivation of enrichment test: We evaluated whether $V_{G_i} - f_i \cdot V_{G_{tot}}$ was statistically significantly positive with a one-sided Z-test, considering that in the null hypothesis of no enrichment, each genomic partition should carry an amount of variance proportional to its number of SNPs. We estimated the variance of the observed enrichment using this equation:

$$Var(V_{G_i} - f_i \cdot V_{G_{tot}}) = Var(V_{G_i}) + f_i^2 \cdot Var(V_{G_{tot}}) - 2 \cdot f_i \cdot Cov(V_{G_i}, V_{G_{tot}}) . \qquad \text{(Eq. 2)}$$

Here, $V_{G_i}$ represents the genetic variance of partition i and $V_{G_{tot}} = \sum_i V_{G_i}$.

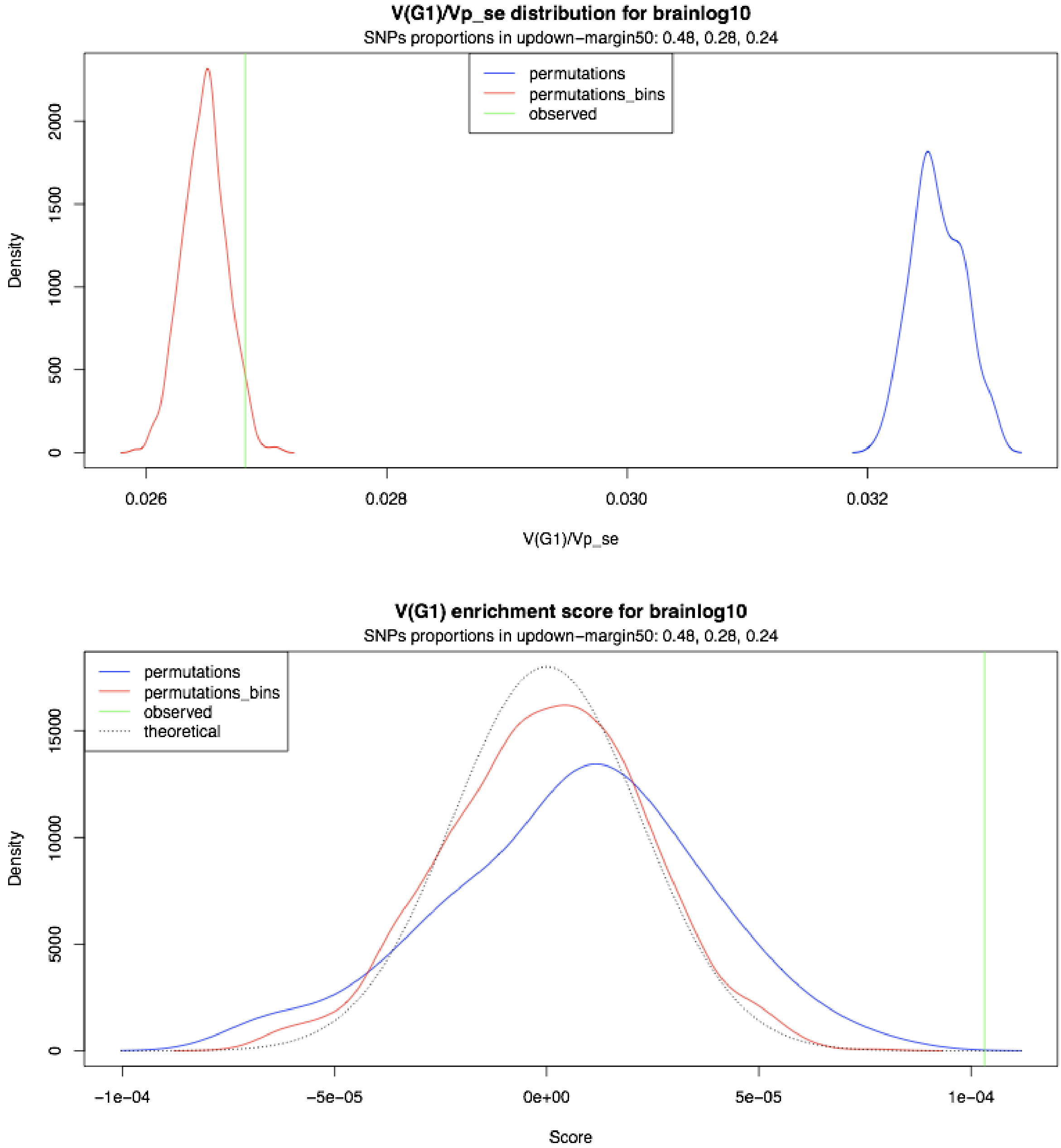


**Figure S1.** Simulation of partitioning the genetic variance between genic, near genic and non genic regions. **Upper panel:** Density of standard errors reported by GCTA in permutation vs standard error reported by GCTA in the original partition. SNPs. **Lower panel:** Density of genic region enrichment for permutations in the null hypothesis of no enrichment computed as the difference between $V_{G_i}$ and $E(V_{G_i}) = V_{G_{tot}} \cdot f_i$. The observed value is the reported value for the real partition. Permutations were made for UK biobank dataset with 1000 iterations. permutations: permutations by SNPs. permutations_bins: permutations by blocks of contiguous SNPs.

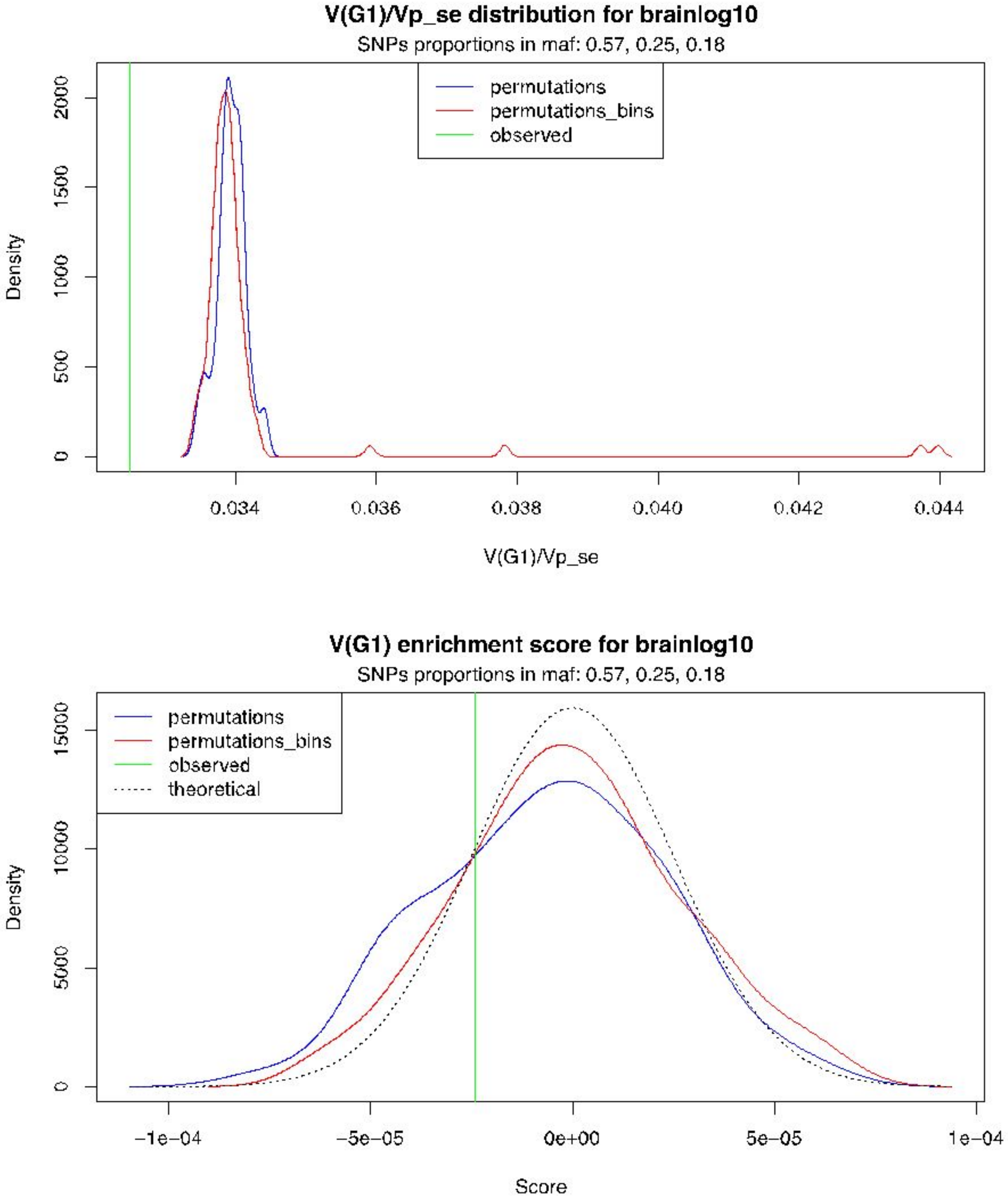


**Figure S2.** Simulation of partitioning the genetic variance in function of MAF (0.05-0.20, 0.20-0.35, 0.35-0.5). **Upper panel:** Density of standard errors reported by GCTA in permutation vs standard error reported by GCTA in the original partition. SNPs. **Lower panel:** Density of low MAF enrichment for permutations in the null hypothesis of no enrichment computed as the difference between $V_{G_i}$ and $E(V_{G_i}) = V_{G_{tot}} \cdot f_i$. The observed value is the reported value for the real partition. Permutations were made for UK biobank dataset with 1000 iterations. permutations: permutations by SNPs. permutations_bins: permutations by blocks of contiguous SNPs.

### 2. Genetic variance estimates in UK Biobank with different set of covariates

We compared the estimations of genetic variance for volumes measured using Freesurfer and FIRST in the UK Biobank project obtained by including or not the 10 first principal components of the genetic relationship matrix, and by including or not brain volume (BV) as a covariate in the linear model. The results with an without the top 10 PCs are included in supplemental table S2.

We observed that $V_G/V_P$ estimates diminished when no covariate was included, meaning that covariates globally capture more environmental variability. The inclusion of brain volume as a covariate did not seem to have a large impact on the estimations.

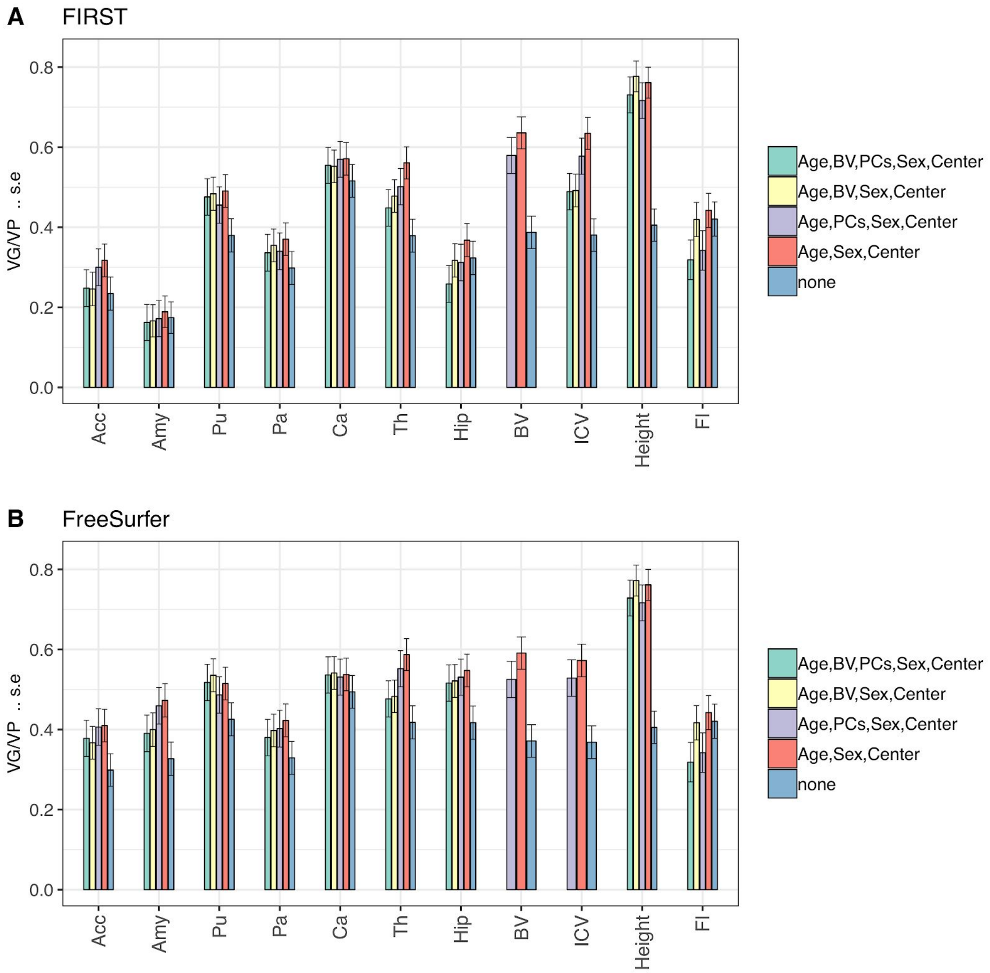


**Figure S3.** Fraction of phenotypic variance (VP) accounted for by genetic variance(VG), computed from all genotyped SNPs for the UK Biobank dataset. $V_G/V_P$ estimates were computed using GCTA REML unconstrained method, for total volumes estimated using either FIRST (A) or FreeSurfer (B). The bar plots also include estimates of $V_G/V_P$ for height and fluid intelligence. The colors represent results obtained with four different sets of covariates (BV denotes the brain volume, PCs the top 10 principal components).

**3. Correlation among the phenotypes studied, and comparison between volume measurement tools.**

We computed the raw correlation matrix of all the phenotypes studied. Figure S3 shows regional volumes computed using Freesurfer, with left and right volumes of each region plot separately. Figure S4 shows the correlation of the volume measurements obtained with Freesurfer and FIRST, with left and right volumes of each structure combined. We observed that patterns of correlations differed across structures, with accumbens and amygdala displaying less correlation than the other structures between their left and right parts. Correlation between volumes measured by either FIRST or FreeSurfer were different across structures, the lowest consistency being found for amygdala (r = 0.621).

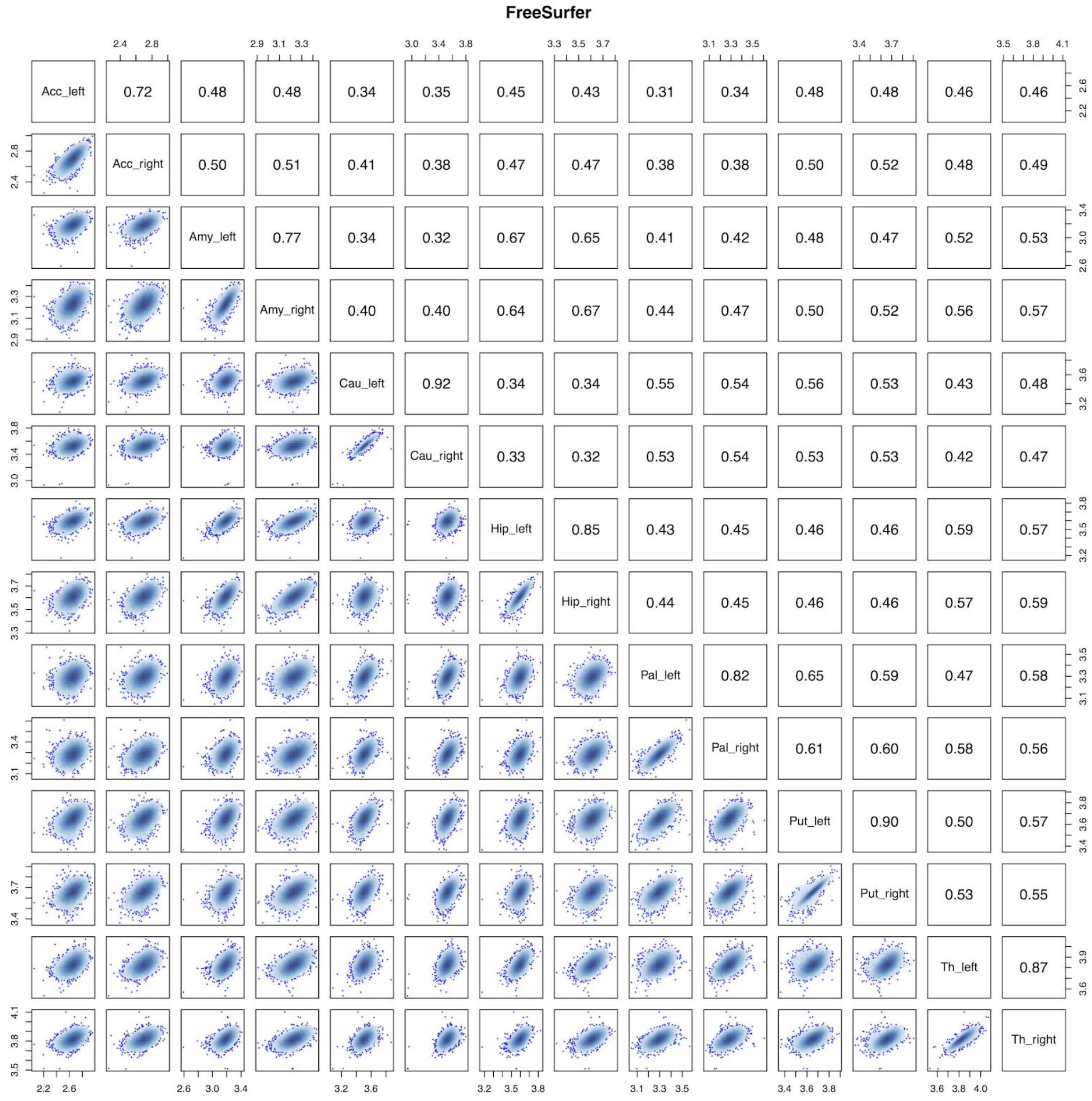


**Figure S4.** Pairs plot of the volumes of the left and right hemispheres of the subcortical structures measured using FreeSurfer (Acc : Accumbens, Amy : Amygdala, Cau : Caudate, Hip : Hippocampus, Pal : Pallidum, Put : Putamen, Th : Thalamus).

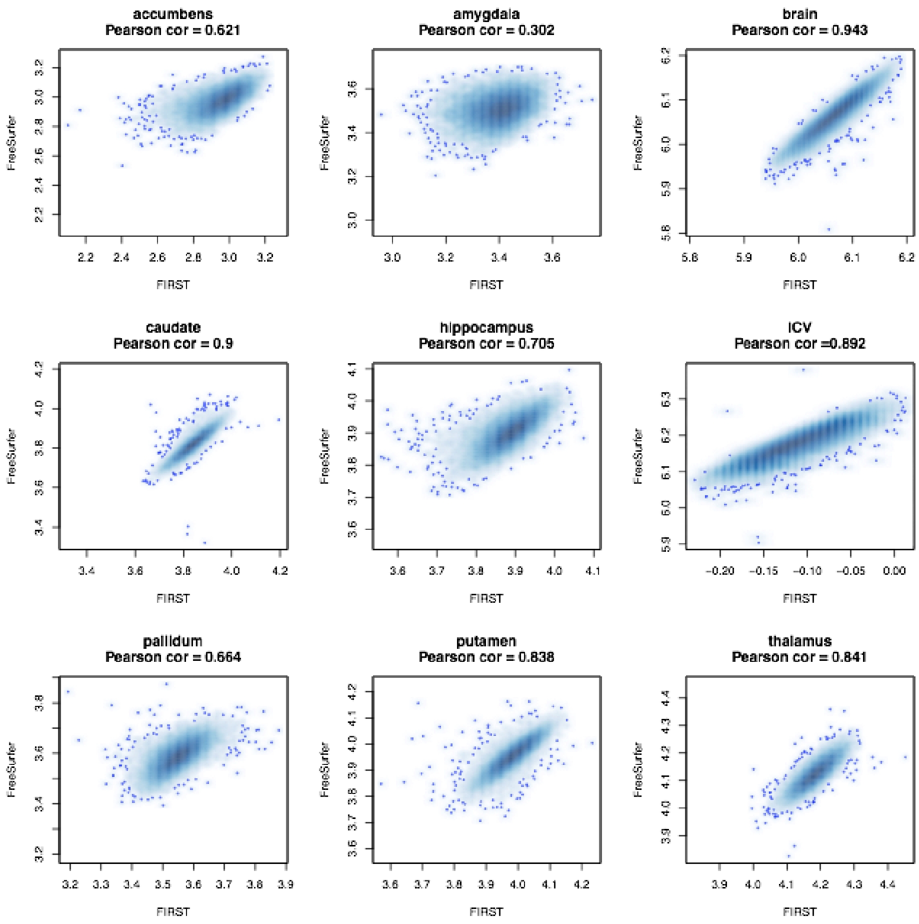


**Figure S5.** Scatter plots of total volumes (sum of left and right hemispheres) measured by FIRST (x-axis) and FreeSurfer (y-axis).

### 4. Comparison of genetic variance estimates with those of previous studies of the UK Biobank and ADNI projects by Elliott et al (2018) and Zhao et al (2018).

Several recent studies have used the UK Biobank data to estimate genetic variance of regional brain volumes. The reported estimates vary across studies, with subcortical volumes, in particular the volume of the amygdala, showing the largest differences. We compared our genetic variance estimates with those obtained by two recent studies (Elliott et al. 2018; Zhao et al. 2018). Zhao et al. (2018) used GCTA to compute $V_G/V_P$ estimates of the left and right volumes of various subcortical structures on a smaller sample of 9,031 subjects from the UK Biobank project. Regional brain volumes were quantified using ANTs (Advanced Normalization Tools software (Avants et al. 2014) together with the MindBoggle-101 atlas (Klein and Tourville 2012). Elliott et al. (2018) reported genetic variance estimates for thousands of phenotypes on 8,411 subjects, using SBAT (Sparse Bayesian Association Test (Elliott et al. 2018)). These phenotypes included the volumes of subcortical structures that we report here. Elliott et al. (2018) provide $V_G/V_P$ estimates obtained from measurements of subcortical structures performed using FSL's FIRST (FMRIB's Integrated Registration and Segmentation Tool (Patenaude et al. 2011)) and FreeSurfer. In previous study of the IMAGEN data we reported only total volumes (left plus right) computed using FIRST. For the sake of comparison, we computed $V_G/V_P$ estimates in the UK Biobank data for the left and right volumes separately, using both FIRST and FreeSurfer (Table S2).

The largest difference of hemispheric variance estimates was observed for the amygdala and the nucleus accumbens (Fig. S6a), which were the smallest structures of those we studied. These structures also displayed the lowest correlation between right and left volumes (Fig. S4) and between FIRST and FreeSurfer measurements (Fig. S5), which suggests that their measurement may be less precise than for larger structures. A similar discrepancy between left and right amygdala volume was observed by both Zhao et al. (2018) and Elliott et al. (2018) (Fig. S6a). Discrepancies between $V_G/V_P$ estimates were more important when comparing different methods to estimate subcortical volumes (FreeSurfer, FIRST, or MindBoggle/ANTs) than when comparing different methods to estimate genetic variance (GCTA or SBAT). The inclusion of the brain volume as a covariate did not change noticeably our variance estimates and thus could not fully explain the differences observed with the study of Zhao et al. (2018) (Fig. S6b). In all cases, however, 95% CIs built from the standard errors of each study contained the estimates of the other studies.

We also compared our $V_G/V_P$ estimates for the ADNI project with those obtained by Zhao et al. (2018). After filtering and merging of the non-imputed genotypes across the two ADNI sub-projects, we included 982 subjects and 210,543 SNPs; Zhao et al. (2018) included 1,023 subjects and 7,368,446 SNPs (after imputation). Despite both analyses using the same covariates, the $V_G/V_P$ estimates showed differences ranging from 7% to 99%. These differences are not surprising, however, given the small sample size, and thus the large standard errors of the estimates (Fig. S6b).

In conclusion, when comparing heritability estimates reported by different studies, the method used to measure volumes had a larger impact on the estimation of genetic variance than the method used for the estimation of heritability. The smallest subcortical structures such as the nucleus accumbens, the amygdala and the pallidum seemed to be the most difficult to segment accurately. Large volume differences both between manual and automated segmentation and between FreeSurfer and FIRST segmentations have been previously reported, in particular for the amygdala (Morey et al. 2009; Schoemaker et al. 2016).

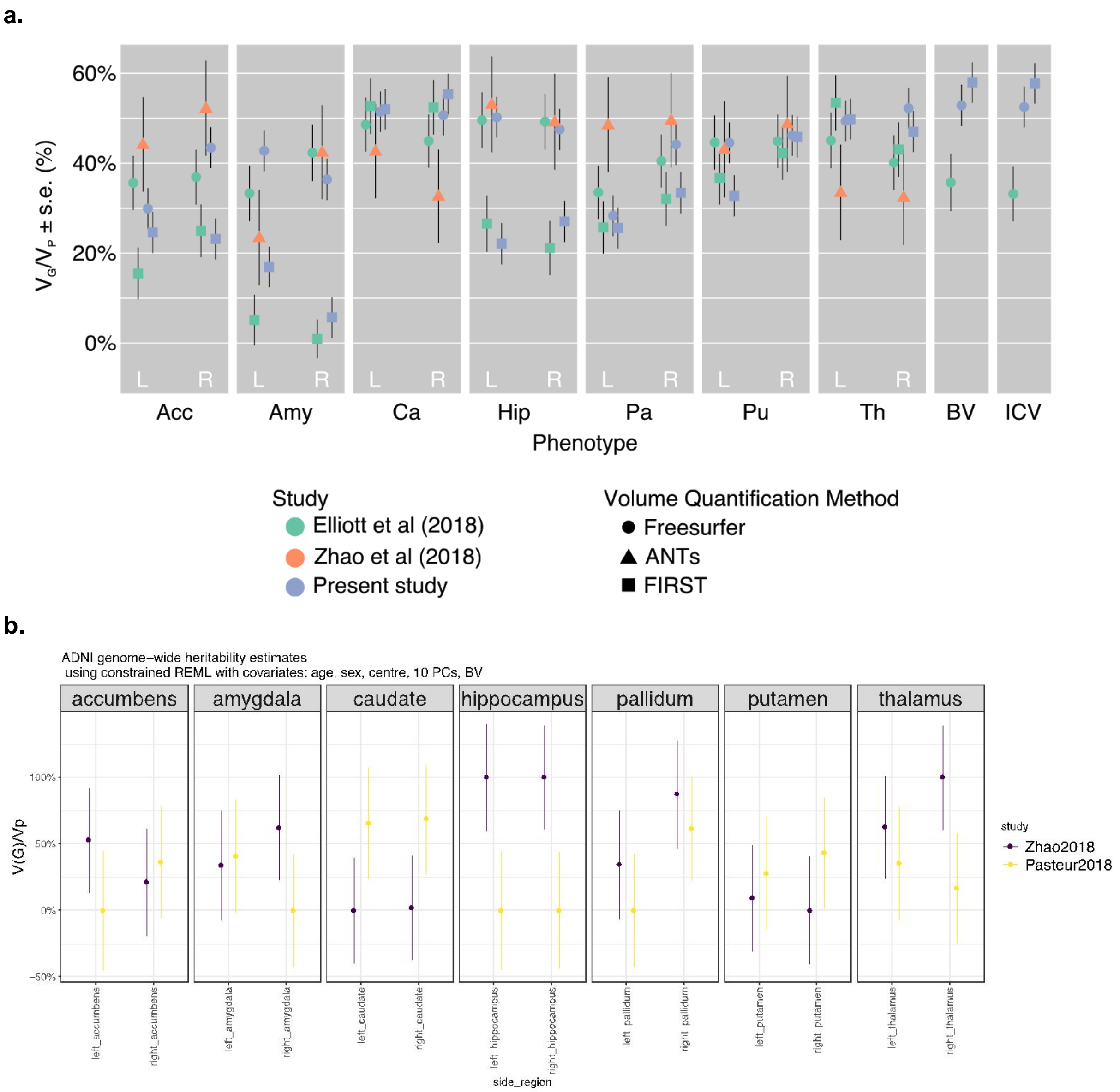


**Figure S6.** (a) Comparison of our estimates of genetic variance ($V_G/V_P$) in the UK Biobank project with two recent studies: Elliot et al. (2018) and Zhao et al. (2018). $V_G/V_P$ estimates were computed for brain, intracranial volume (ICV), and subcortical volumes of the right and left hemispheres measured using different methods (FreeSurfer, or MindBoggle/ANTs, FIRST) represented by the shapes of the dots. In yellow are represented the estimates of $V_G/V_P$ we computed using GCTA REML unconstrained method, the other $V_G/V_P$ estimates were published by Elliott et al. 2018 (green) and Zhao et al. 2018 (orange). Age, sex, center, and the top 10 principal components were included as covariates. The error bars show the standard errors of the $V_G/V_P$ estimates. (b) Fraction of phenotypic variance ($V_P$) accounted for by genetic variance ($V_G$), computed from all genotyped SNPs for the volumes of six subcortical structures in the ADNI dataset. We observed that the sample size was too small to get reliable estimations. Estimates published by Zhao et al., 2018 are shown in violet, estimates we obtained using GCTA REML constrained method with covariates similar to the ones used by Zhao et al. (age, sex, center, 10 top principal components, brain volume, ADNI phase) are shown in yellow.

### 5. Comparison of phenotypic and genetic correlations

We computed matrices of phenotypic correlations ($r_P$ in Fig. S7, left panel) and genetic correlations ($r_G$ in Fig. S7, right panel). The correlations among pairs of phenotypes were computed using GCTA. We used Ward hierarchical clustering based on euclidean distance to cluster the phenotypes.

Additionally, we computed environmental correlations which are the environmental counterpart of genetic correlations (Fig. S8, left panel). They represent the correlation between the environmental effects of two traits. They were computed using the environmental variances of each trait and the environmental covariance between the two traits output by GCTA ( $r_E = \frac{Cov(E_{tr1}, E_{tr2})}{\sqrt{Var(E_{tr1}) \cdot Var(E_{tr2})}}$ ). Standard errors were computed with the delta method. The concordance between genetic and environmental correlation was medium ($R^2 = 0.47$) (Fig S8, right panel).

Correlations were overall strong and significant, especially between the left and right parts of the same structure. Figure S9 shows the values of the differences between genetic and environmental correlations ($r_G$ - $r_E$, left panel) and genetic and phenotypic ($r_G$ - $r_P$, right panel) are shown in the lower triangular part of the matrix. Differences between genetic and environmental correlations were larger than the differences between genetic and phenotypic correlations. Statistical significance was assessed using a Z-test to determine whether the differences were non null (Benjamini Yekutieli FDR adjusted *P<0.05, **P<0.01, ***P<0.001). Table S5 provides the values of the genetic, phenotypic, and environmental correlations and the results of their comparison.

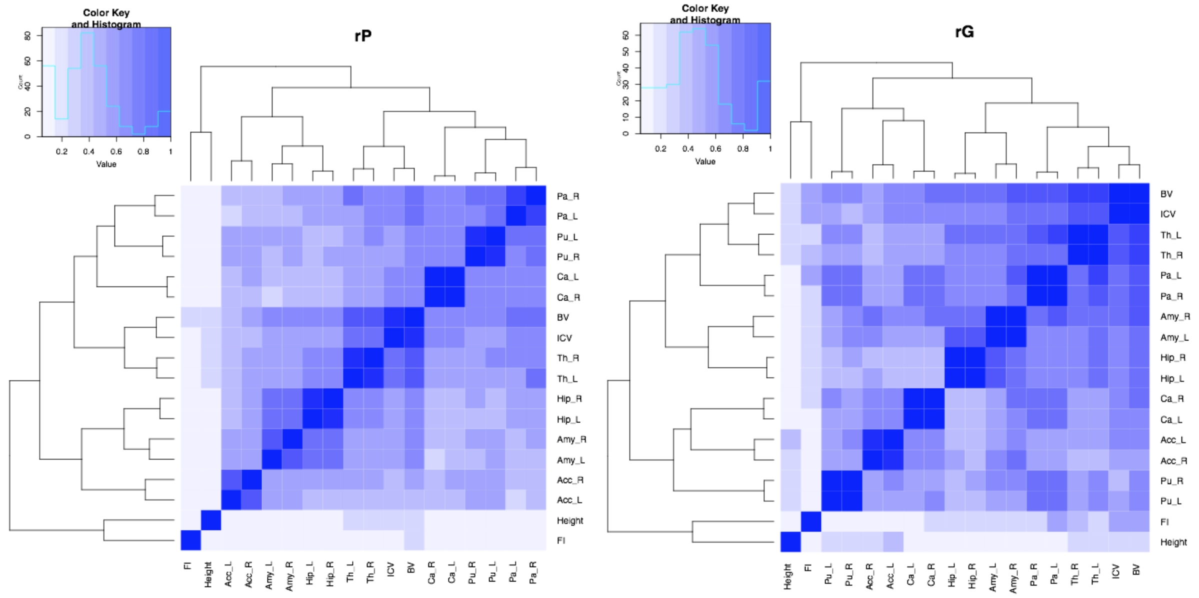


**Figure S7**. Heat maps of phenotypic correlations ($r_P$, left panel) and genetic correlation ($r_G$, right panel) values computed with GCTA. The following phenotypes are represented: the left and right hemisphere volumes of accumbens (Acc_L and Acc_R), amygdala (Amy_L and Amy_R), caudate (Cau_L and Cau_R), hippocampus (Hip_L and Hip_R), pallidum (Pa_L and Pa_R), putamen (Pu_L and Pu_R), and thalamus (Th_L and Th_R) ; brain volume (BV), intracranial volume (ICV), height and fluid intelligence (FI) are also represented. Ward hierarchical clustering based on euclidean distance was used to cluster the phenotypes.

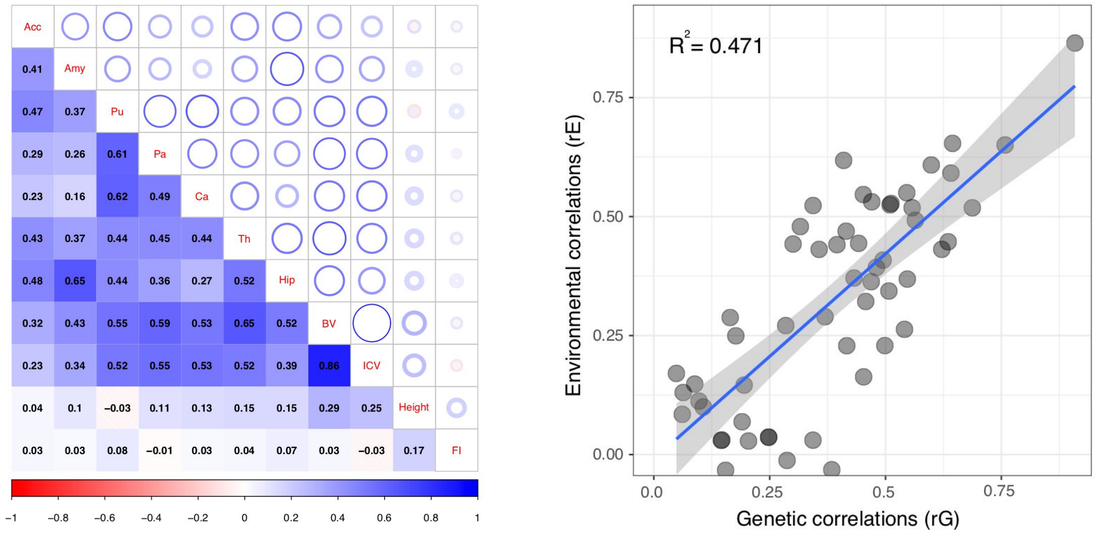


**Figure S8.** Left: values of environmental correlations are shown in the lower triangular part of the matrix. Circles's diameter is proportional to the strength of the correlation, and a thickness proportional to the size of the confidence interval. Right: scatter plot of environmental versus genetic correlations. Genetic and environmental correlations were adjusted for age, center, sex, and the top 10 principal components of the genetic relationship matrix.

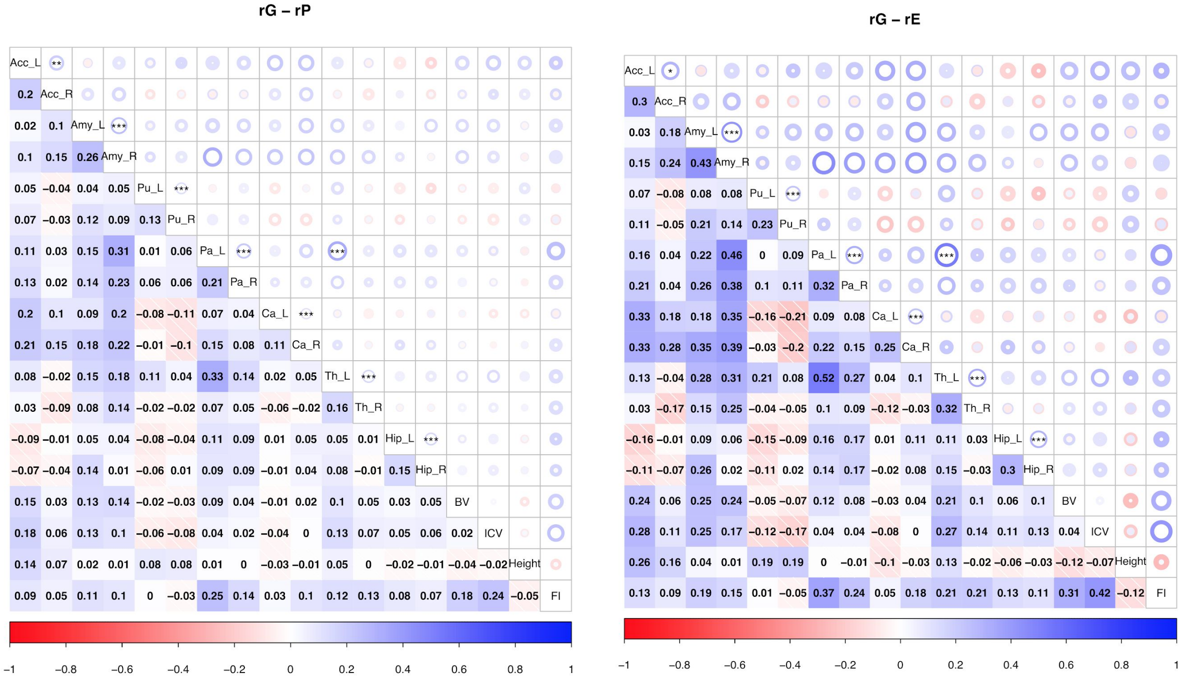


**Figure S9**. Values of the differences between genetic and phenotypic correlations ($r_G$ - $r_P$, left) and genetic and environmental correlations ($r_G$ - $r_E$, right) are shown in the lower triangular part of the matrix. Circles' diameter is proportional to the difference and a thickness proportional to the size of the confidence interval. Stars indicate statistical significance from a Z-test testing if the differences are non null (Benjamini Yekutieli FDR adjusted *$P$<0.05, **$P$<0.01, ***$P$<0.001). The following phenotypes are represented: the left and right hemisphere volumes of accumbens (Acc_L and Acc_R), amygdala (Amy_L and Amy_R), caudate (Cau_L and Cau_R), hippocampus (Hip_L and Hip_R), pallidum (Pa_L and Pa_R), putamen (Pu_L and Pu_R), and thalamus (Th_L and Th_R) ; brain volume (BV), intracranial volume (ICV), height and fluid intelligence (FI) are also represented.

**Accounting for the effect of phenotypic measurement error**

Let x and y be two measured phenotypes, they can be decomposed into their real values $x'$ and $y'$ and their measurement errors $\varepsilon_x$ and $\varepsilon_y$ : $x = x' + \varepsilon_x$ and $y = y' + \varepsilon_y$ .

The correlation between the two measured phenotypes can be expressed like this:

$$r_{x,y} = \frac{cov(x, y)}{\sqrt{var(x)\, var(y)}} = \frac{cov(x'+\varepsilon_x, y'+\varepsilon_y)}{\sqrt{var(x'+\varepsilon_x)\, var(y'+\varepsilon_y)}} \qquad \text{(Eq. 3)}$$

Considering that there is no correlation between measurement errors and the real values, one can develop the following formula:

$$r_{x,y} = \frac{cov(x', y')+cov(\varepsilon_x, \varepsilon_y)}{\sqrt{(var(x')+var(\varepsilon_x))(var(y')+var(\varepsilon_y))}}$$

$$= \frac{cov(x', y')}{\sqrt{var(x')var(y')}} \cdot \frac{cov(x', y')+cov(\varepsilon_x, \varepsilon_y)}{cov(x', y')} \cdot \sqrt{\frac{var(x')}{var(x')+var(\varepsilon_x)}} \cdot \sqrt{\frac{var(y')}{(var(y')+var(\varepsilon_y))}} \qquad \text{(Eq. 4)}$$

$\frac{var(x')}{var(x')+var(\varepsilon_x)}$ and $\frac{var(y')}{var(y')+var(\varepsilon_y)}$ can be estimated as being the intraclass correlation coefficient (ICC) respectively for *x* and *y* (called $r_x$ and $r_y$).

We used a generalised version of the intraclass correlation coefficient to estimate $\frac{cov(x', y')}{cov(x', y')+cov(\varepsilon_x, \varepsilon_y)}$ (called $r_c$): When *x* and *y* are measured for *N* different subjects with *k* repeated measurements, $r_c = \frac{1}{k(k-1)Nc} \sum_{i=1}^{k} \sum_{j=1, j \neq i}^{k} \sum_{n=1}^{N} (x_{n,i} - \bar{x})(y_{n,j} - \bar{y})$ where $\bar{x}$ and $\bar{y}$ are the respective means for all *x* and *y* measurements, and $c = \frac{1}{kN} \sum_{i=1}^{k} \sum_{n=1}^{N} (x_{n,i} - \bar{x})(y_{n,i} - \bar{y})$ .

We were thus able to obtain an estimation of the phenotypic correlation adjusted for measurement errors:

$$r_{x',y'} = r_{x,y} \frac{r_c}{\sqrt{r_x r_y}} \qquad \text{(Eq. 5).}$$

Values of $r_{x',y'}$ are available in the second sheet of Table S5, values of $r_x$ , $r_y$ , and $r_c$ are available in the third sheet of Table S5. The estimates of $V_G/V_P$ adjusted for measurement errors using the ICC values of each phenotype are shown in Figure S10.

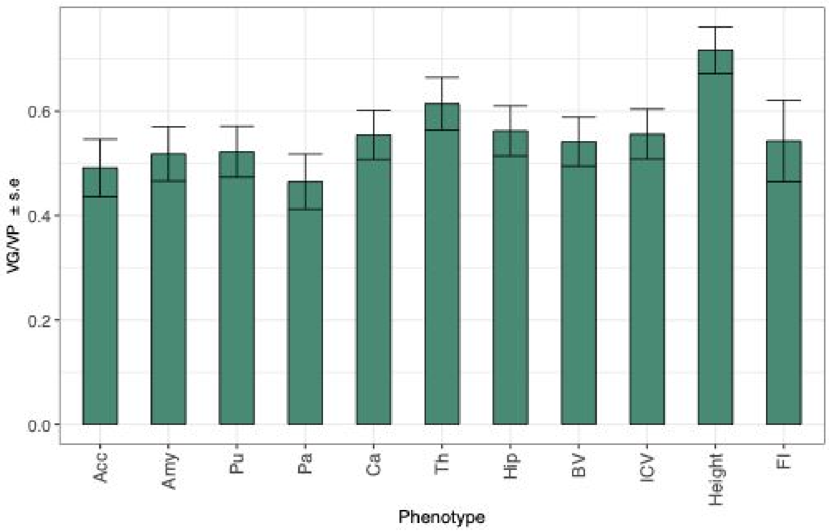


**Figure S10.** Adjusted proportion of variance captured by common genotyped variants ($V_G/V_P$) for brain regions, height and intelligence score.

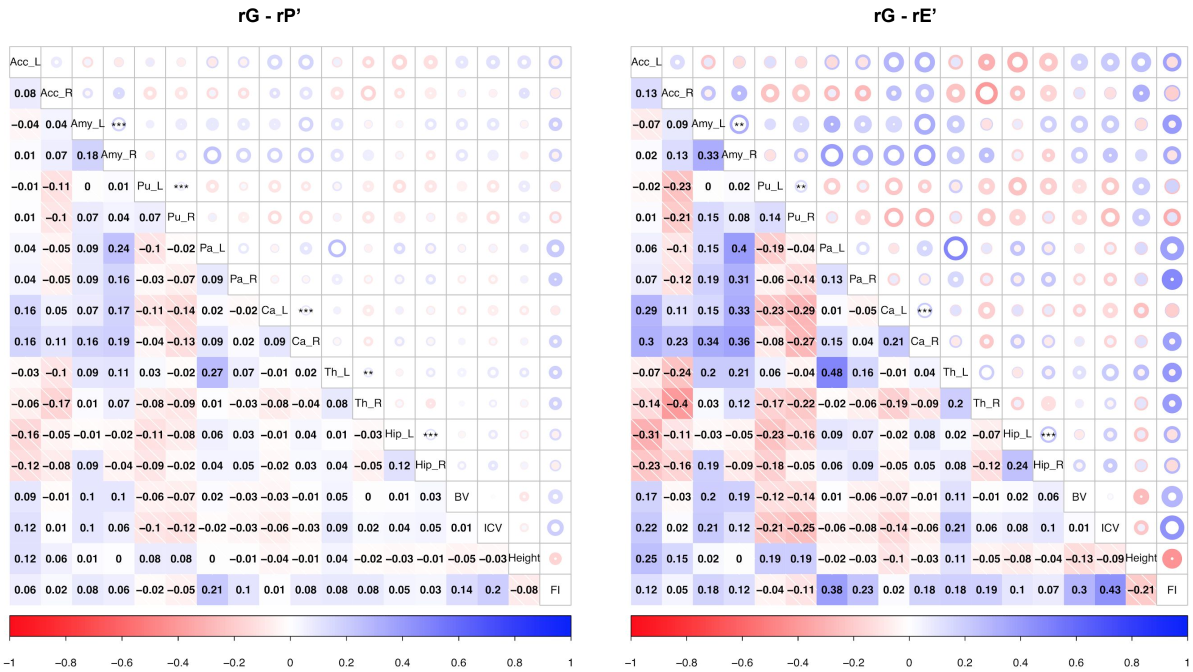


**Figure S11**. Values of the differences between genetic and corrected phenotypic correlations ($r_G$ - $r_P$', left panel), and between genetic and corrected environmental correlations ($r_G$ - $r_E$', right panel) are shown in the lower triangular part of the matrix. Circles' diameter is proportional to the difference and a thickness proportional to the size of the confidence interval. Stars indicate statistical significance from a Z-test testing if the differences are non null (Benjamini Yekutieli FDR adjusted **P*<0.05, ***P*<0.01, ****P*<0.001). The following phenotypes are represented: the left and right hemisphere volumes of accumbens (Acc_L and Acc_R), amygdala (Amy_L and Amy_R), caudate (Cau_L and Cau_R), hippocampus (Hip_L and Hip_R), pallidum (Pa_L and Pa_R), putamen (Pu_L and Pu_R), and thalamus (Th_L and Th_R) ; brain volume (BV), intracranial volume (ICV), height and fluid intelligence (FI) are also represented.

### 6. Distribution of $V_G/V_P$ estimates for simulated phenotypes for different sample sizes

We studied how the distribution of heritability estimates differed from their theoretical distribution as a function of sample size. $V_G/V_P$ estimates were obtained from 1,000 simulated phenotypes with a heritability of $V_G/V_P$ = 50% for sub-samples of N = 800, N = 400, N = 200 and N = 100 subjects.
Figure S12 show simulations based on the ADNI project and Figure S13 show simulations based on the UK Biobank project.

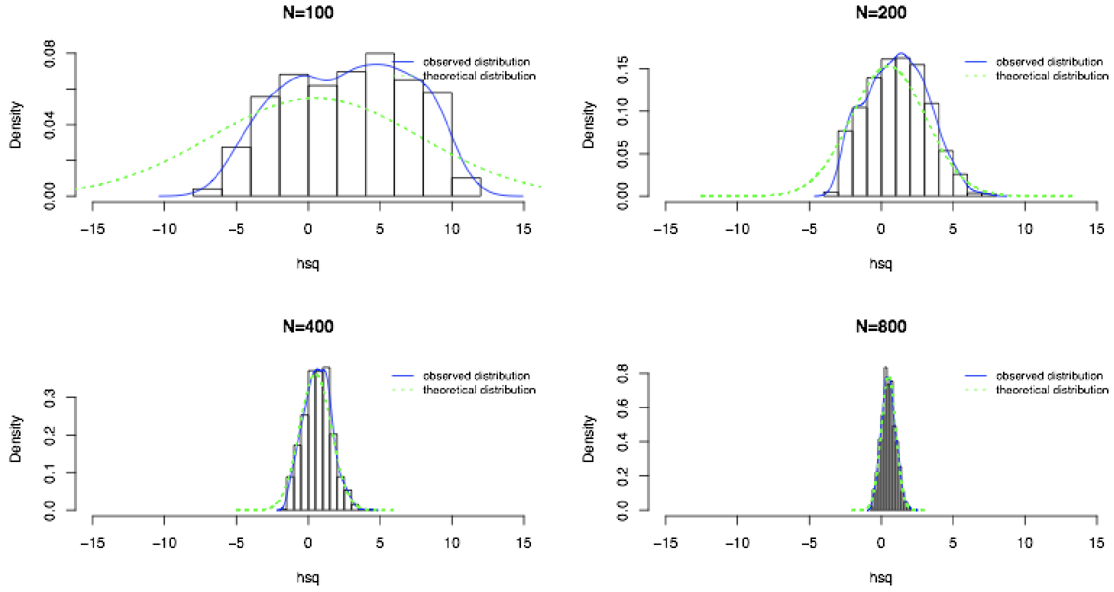


**Figure S12.** Distribution of estimated values for $V_G/V_P$ of simulated phenotypes for different sub-sample sizes in ADNI dataset, compared with the expected (theoretical) distribution.

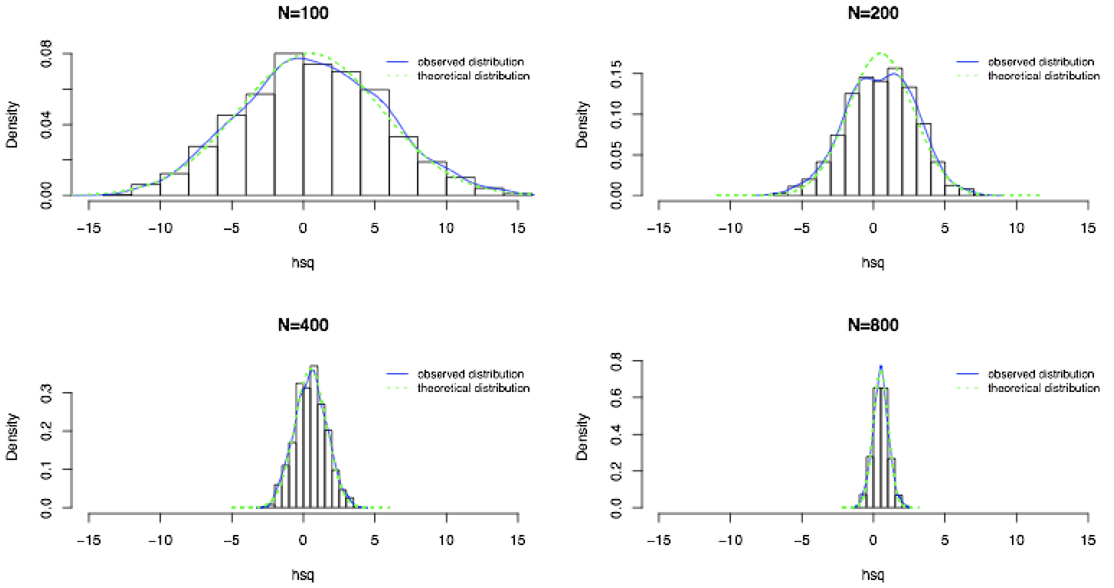


**Figure S13.** Distribution of estimated values for $V_G/V_P$ of simulated phenotypes for different sub-sample sizes in UK Biobank dataset, compared with the expected (theoretical) distribution.

#### 7. Comparison between meta-analysis and mega-analysis

Fig. S14 shows the difference in standard errors obtained when a single group is analysed (labelled as "merging") versus a meta-analytic approach combining several small groups (labelled as "meta-analysis"). The plot represents the change in standard error after including an additional dataset for GREML analysis.

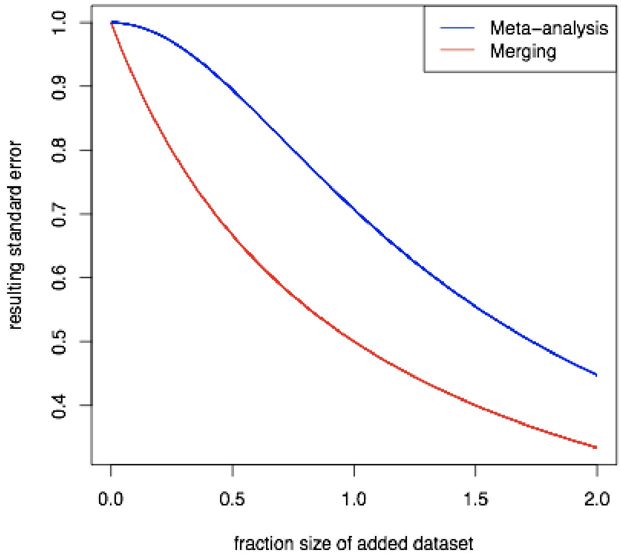


**Figure S14.** Resulting standard error after addition of another dataset for GREML analysis. The x-axis represents relative size of the added dataset compared with the initial dataset. The y-axis represents relative standard error of the combinated estimate compared with the standard error obtained only with the first dataset. Meta-analysis: resulting standard error computed as $\frac{1}{\sqrt{1+fraction^2}}$. Merging: resulting standard error computed as $\frac{1}{1+fraction}$.

### 8. Genome-wide polygenic scores

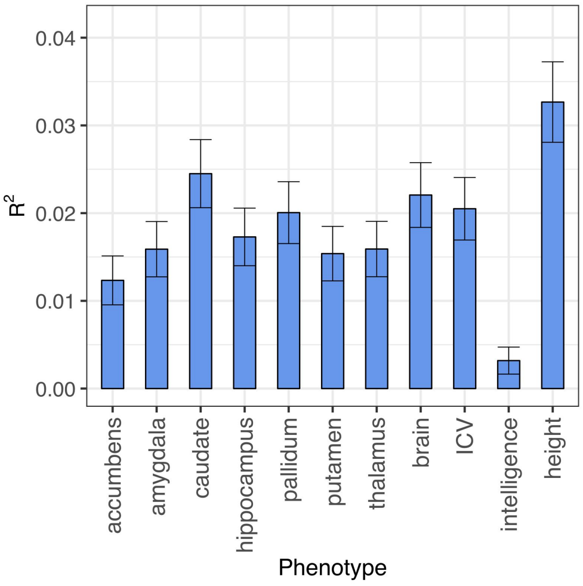


**Figure S15.** Variance captured by genome-wide polygenic scores (coefficient of determination $R^2$) for each phenotype. The analyses were performed on the residuals of the linear regression including the effects of age, sex, imaging centre, and the 10 first principal components of the GRM as covariates. Error bars represent the 95% confidence intervals.

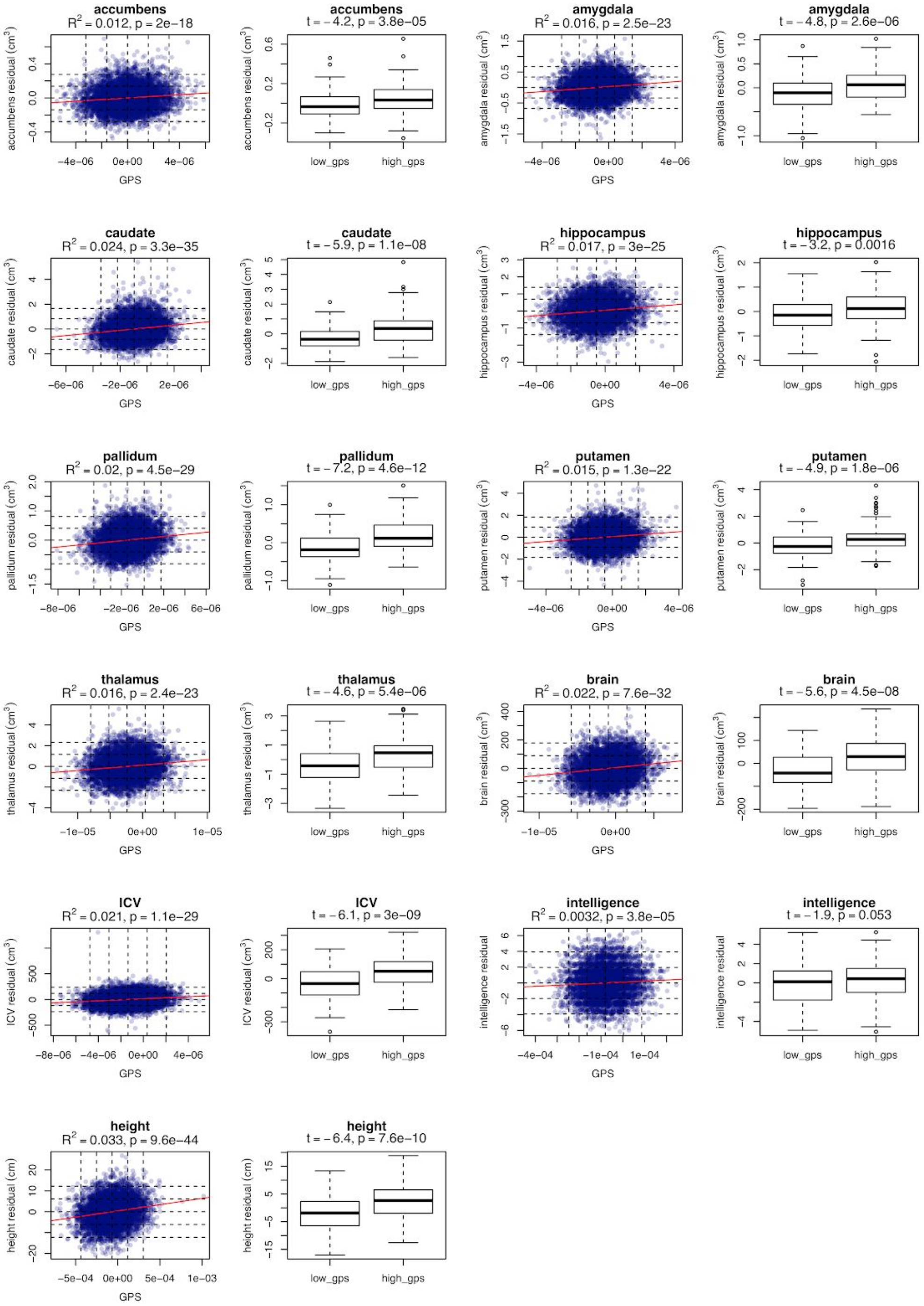


**Figure S16.** For each phenotype**,** two plots are shown: a scatter plot of the phenotype residuals (after covarying by age, sex, imaging centre and genetic principal components) as a function of the genome-wide polygenic scores (GPS), and boxplots showing the distribution of the phenotype residuals of the individuals located more than minus (low_gps) or plus (high_gps) two standard deviations away from the average GPS. Units are $cm^3$ for volumes and cm for height. The variance captured by genome-wide polygenic scores (coefficient of determination $R^2$) and the p-value of the Fisher test testing whether $R^2$ is different from zero are given for the scatter plots. The t-value and p-value of the Student test comparing the individuals with low and high GPS are given for the boxplots.

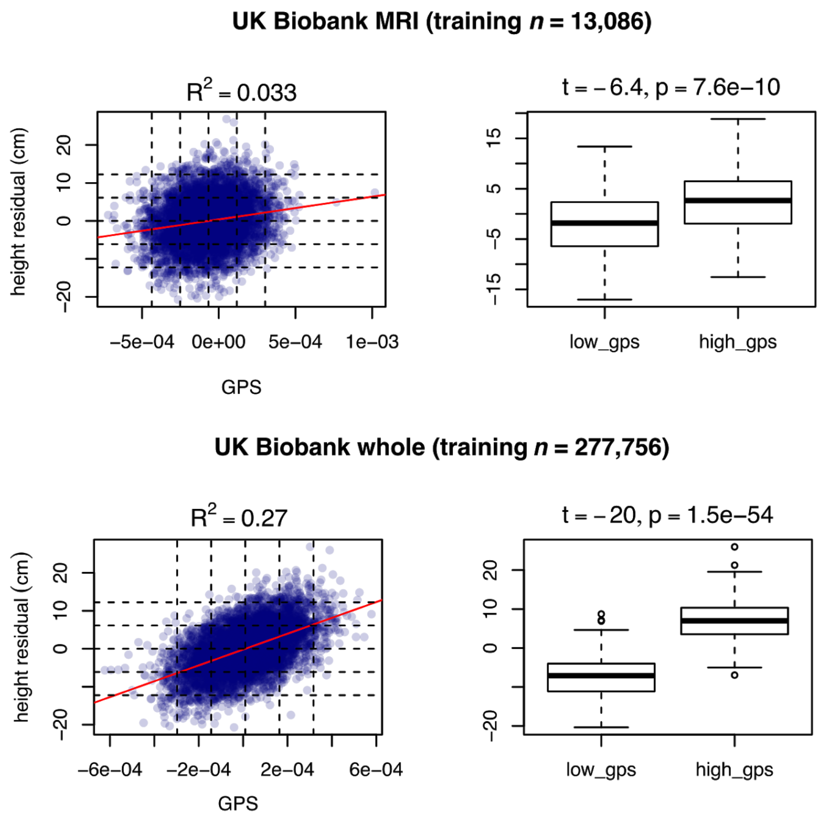


**Figure S17.** Comparison of the prediction of height with genome-wide polygenic scores using as base dataset the MRI cohort of UK Biobank (n = 13,086, top panel) and the whole UK Biobank cohort without subjects included in the target dataset (n = 277,756, bottom panel). The additional MRI cohort of UK Biobank (n = 6,189) was used as target dataset.

For each base dataset**,** two plots are shown: a scatter plot of the phenotype residuals (after covarying by age, sex, imaging centre and genetic principal components) as a function of the genome-wide polygenic scores (GPS), and boxplots showing the distribution of the phenotype residuals of the individuals located more than minus (low_gps) or plus (high_gps) two standard deviations away from the average GPS. Units are $cm^3$ for volumes and cm for height. The variance captured by genome-wide polygenic scores (coefficient of determination $R^2$) are given for the scatter plots. The t-value and p-value of the Student test comparing the individuals with low and high GPS are given for the boxplots.

### 9. Supplementary Table Legends

**Table S1: SNP-heritability results.**

Sheet1: SNP-heritability computed with the GCTA GREML method (including age, sex, centre and the top 10 PCs of the GRM as covariates) for each of the studied phenotypes in UK Biobank and five ENIGMA cohorts.

Sheet2: SNP-heritability estimates obtained by averaging across the six cohorts using the inverse variance-weighted average method.

**Table S2:** SNP-heritability computed with GCTA GREML for UK Biobank (including age, sex, centre), with (Table S2.1) and without (Table S2.2) including the top 10 principal components of the GRM matrix as covariates.

**Table S3: SNP-heritability and enrichment results for several partitions using the UK Biobank project data.**

Table S3.1 provides data on four partitions based on genic status. The first set includes all SNPs within the 66,632 RefSeq gene boundaries of the hg19 assembly ("genic-margin0"), the two further sets include also SNPs 0 to 20 kbp ("updown-margin20") and 20kbp to 50 kbp upstream and downstream of each gene ("updown-margin20-50"), and a last set includes the SNPs not located in regions less than 50 kbp upstream or downstream of genes (nongenic-margin50).

Table S3.2 provides data on four partitions based on minimum allele frequency (MAF): from 0.1 to 5%, from 5 to 20%, from 20 to 35% and from 35 to 50%.

Table S3.3 provides data on three partitions (genic in gene set region, genic not in gene set region, non-genic) based on preferential central nervous system (CNS) gene expression (Raychaudhuri et al. 2010; Lee, DeCandia, et al. 2012) using ± 50 kbp as gene boundaries.

Table S3.4 provides data on three partitions (genic in gene set region, genic not in gene set region, non-genic) based on markers of brain cell types from the scRNA-Seq study of Skene et al. (2018) (PMID: 29785013). We used the specificity metric, provided by the authors in R datafiles at http://www.hjerling-leffler-lab.org/data/scz_singlecell/, for each gene across 24 brain cell types. In order to obtain a partition with a sufficient number of SNPs, we built a gene set (using ± 20 kbp as gene boundaries) of brain cell types markers by taking the union of the genes with the top 1% specificity scores for each cell type.

Table S3.5 provides data on three partitions (genic in gene set region, genic not in gene set region, non-genic) based on markers of brain cell types from the scRNA-Seq study of Li et al. (2018) (PMID: 30545854). The union of the genes in the Table S6 of the manuscript were used to build the gene set (using ± 20 kbp as gene boundaries).

Table S3.6 provides data on three partitions (genic in gene set region, genic not in gene set region, non-genic) based on markers of different time periods of human brain development from the scRNA-Seq study of Li et al. (2018) (PMID: 30545854). The union of the genes in the Table S7 of the manuscript were used to build the gene set (using ± 20 kbp as gene boundaries).

For each partition i, enrichment was computed using a Z-score comparing the estimated genetic variance VGi of the SNP set i to the SNP-set genetic variance fi*VGi expected under no enrichment, the corresponding enrichment p-value comparing this Z-score to zero with a Z-test, the number of individuals, the p-value of the likelihood ratio test output by GCTA.

**Table S4. Meta-Analysis of the genetic partitioning of SNP-heritability across the six datasets.**
The partitioning of SNP-heritability estimates were averaged across the six datasets using the inverse variance-weighted average method.
Table S4.1 provides four partitions based on genic status.
Table S4.2 provides four partitions based on minimum allele frequency (MAF).
Table S4.3 provides two partitions based on preferential central nervous system (CNS) gene expression (Raychaudhuri et al. 2010; Lee, DeCandia, et al. 2012) using ± 50 kbp as gene boundaries.
For each partition i, the following results are provided: $f_i$ the average fraction of SNPs belonging to the SNP set i, the average values of $V_{Gi}/V_P$ and $V_{Gi}/V_G/f_i$ and the associated standard errors, the enrichment p-value of the Z-test comparing $V_{Gi}/V_G/f_i$ to 1.

**Table S5: Genetic, phenotypic, and environmental correlations in UK Biobank.**
Table S5.1 provides the values of the genetic, phenotypic, and environmental correlations, together with their standard errors. Differences between $r_G$ and $r_E$ (column dr_$r_G r_E$), and $r_G$ and $r_P$ (column dr_$r_G r_P$) were computed and their associated standard errors (column dr_$r_G r_E$_SE and column dr_$r_G r_P$_SE) was estimated using the delta method using the output of the GCTA REML bivariate method. A Z-test was used to compare these differences to zero, the p-values are available in columns p.dr_$r_G r_E$ and p.dr_$r_G r_P$. The column nind contains the number of individuals available with both annotated phenotypes.
Table S5.2 provides the same values as for Table S5.1 but for phenotypic and environmental correlations values that were corrected for measurement error (as described in the 5. of this supplementary document).
Table S5.3 provides the values used to correct the phenotypic correlations for measurement errors (see equation Eq. 5 of this supplementary document).

**Table S6:** Distribution of estimated values for $V_G/V_P$ of simulated phenotypes for different sub-sample sizes in ADNI dataset. For four sample sizes, 1000 sub-sampling were performed without replacement. nb_ind: number of sub-sampled individuals, nb_conv: number of times GREML converged and an estimate was given, th_mean: theoretical mean for estimates (equal to the simulated $V_G/V_P$), obs_mean, obs_mean_se: observed mean of estimates and standard error, p_mean: p value for difference between theoretical and observed mean (z-test), th_sd: theoretical standard deviation of estimates (computed as the mean of standard errors reported by GCTA), obs_sd, obs_sd_se: observed standard deviation of estimates and standard error, p_sd: p value for difference between theoretical and observed standard deviation (z-test), D_ks, p_ks: effect size and p-value of Kolmogorov-Smirnov test of concordance between theoretical normal distribution and observed distribution.

**Table S7:** Distribution of estimated values for $V_G/V_P$ of simulated phenotypes for different sub-sample sizes in UK Biobank dataset.

**Table S8:** Proportion of captured phenotypic variance by genome-wide polygenic scores and covariates. mean_pheno: mean value of the raw phenotype, sd_pheno: standard deviation of the raw phenotype, Rsq_age_only: proportion of the raw phenotype captured by age, Rsq_pca_only: proportion of the raw phenotype captured by genetic principal components, Rsq_sex_only: proportion of the raw phenotype captured by sex, Rsq_centre_only: proportion of the raw phenotype captured by imaging centre, Rsq_gps_only: proportion of the raw phenotype captured by genome-wide polygenic score, sd_resid: standard deviation of residual phenotype after covarying by age, pca, sex and centre, Rsq: proportion of residual phenotype captured by polygenic score, SErsq: standard error of $R^2$, LCL: lower 95% confidence interval limit, UCL: upper 95% confidence interval limit.
